# Energetic driving force for LHCII clustering in plant membranes

**DOI:** 10.1101/2023.04.19.537479

**Authors:** Premashis Manna, Madeline Hoffmann, Thomas Davies, Katherine H. Richardson, Matthew P. Johnson, Gabriela S. Schlau-Cohen

## Abstract

Plants protect themselves against photodamage from excess energy using a process known as non-photochemical quenching (NPQ). A significant fraction of NPQ is induced by a ΔpH across the membrane, which changes the conformation, composition, and organization of the antenna complexes. In particular, clustering of the major light-harvesting complex (LHCII) has been observed, yet the thermodynamic driving force behind this reorganization has not been determined, largely because measurements of membrane protein interaction energies have not been possible. Here, we introduce a method to quantify membrane protein interaction energies and its application to the thermodynamics of LHCII clusters. By combining single-molecule measurements of LHCII-proteoliposomes at different protein densities and a rigorous analysis of LHCII clusters and photophysics, we quantified the LHCII-LHCII interaction energy to be approximately -5 *k*_*B*_*T* at neutral pH and at least -7 *k*_*B*_*T* at acidic pH. From these values, we found the thermodynamic driving force for LHCII clustering was dominated by these enthalpic contributions. Collectively, this work captures the membrane protein-protein interactions responsible for LHCII clustering from the perspective of equilibrium statistical thermodynamics, which has a long and rich tradition in biology.

## INTRODUCTION

Biology is replete with protein-protein interactions that change the enthalpy (H) and entropy (S) of the system. According to the Second Law of Thermodynamics, for these interactions to form spontaneously, they must lead to an increase in total entropy (S). Stated differently, a spontaneous process inevitably results in a decrease in the state function called the Gibbs free energy (G), defined as G=H-TS, where T is the temperature. The use of ideas from equilibrium thermodynamics and its microscopic partner, equilibrium statistical mechanics, has a long and rich tradition in biology^1,2^. In particular, the application of statistical thermodynamics to protein-protein interactions has been a powerful tool for understanding processes such as multivalent binding^3^, complex formation^4^, and hydration of protein complexes^5^. However, these studies have been limited to globular proteins because of the challenges associated with studies of membrane proteins and so a thermodynamic understanding of the many biological processes that occur within membranes is missing. The most abundant membrane protein on Earth is light-harvesting complex II (LHCII), the primary antenna protein in green plants. Despite its importance, only a few studies have addressed the statistical thermodynamic aspects of photosynthetic light harvesting, and such studies have been limited to the properties of individual proteins^6,7^.

In the first steps of photosynthesis, solar energy is absorbed by a network of LHCII and other homologous antenna proteins and used to drive a cascade of biochemical processes that ultimately convert CO_2_ to carbohydrates. Under high light, such as direct sun, the photosynthetic machinery is susceptible to damage from over-excitation and the resultant generation of harmful singlet oxygen^8^. To cope with this, plants have evolved photoprotective mechanisms known as non-photochemical quenching (NPQ)^9,10^. Intense research over the last few decades resolved aspects of NPQ, including the sites, triggers, and quenchers^8^. In the fast, energy-dependent component of NPQ, termed qE, excess energy is dissipated as heat. qE is activated by a ΔpH across the membrane, which is generated by proton accumulation from water splitting^11^. Up to 60% of qE occurs in LHCII^12^, where conformational changes of the protein activate dissipative pathways within the embedded chlorophyll (Chl) and carotenoids (Car). An equilibrium between the light-harvesting and dissipative conformations of LHCII is thought to regulate the extent of qE in plants and algae^13^. The free energy differences that describe this equilibrium have been studied for individual LHCII under extreme temperature or pressure conditions^6,7^. However, the high protein density (80%) of the thylakoid membrane means that in vivo LHCII functions in combination with surrounding proteins. Numerous studies have shown that LHCII has a strong tendency to form clusters^14–19^, particularly under qE conditions^20^, despite the net negative charge of -21e for LHCII trimers^21^. The thermodynamic origin of the clustering in the face of electrostatic repulsion is missing in the literature, including the related parameters, *e*.*g*., entropy, enthalpy, and Gibbs free energy.

In this article, we introduce a method to quantify the interaction energies of LHCII and the thermodynamic driving forces associated with membrane reorganizations. We employed single-molecule spectroscopy of LHCII-proteoliposome samples and modeled the photophysics and thermodynamics of LHCII clusters within the proteoliposomes, which were fit to the experimental single-molecule data to extract the LHCII-LHCII interaction energies. The interaction energy was found to be attractive in nature, consistent with observations of clustering, and in the range of several *k*_*B*_*T* s, where *k*_*B*_ is the Boltzmann constant. The interaction energy at low pH increased in magnitude by *∼*30% compared to neutral pH, which drives a transition from moderately clustered LHCII configurations to strongly clustered ones. Quantification of the free energy change using the measured interaction energies established that the pH-driven clustering of LHCII is primarily an enthalpy-driven process. The enthalpy of the cluster formation is in the tens of *k*_*B*_T (*i*.*e*., few kJ/mol), indicating a driving force strong enough to induce LHCII clustering yet weak enough force for easy reversibility between configurations.

## RESULTS

### LHCII proteoliposome system to probe protein-protein interactions

We investigated the protein-protein interactions between LHCII using proteoliposomes with a systematically increasing number of LHCII. The proteoliposomes were prepared by first extruding the thylakoid lipid mixture through a polycarbonate membrane with 25 nm pore radii to generate liposomes of a well-controlled size. The liposomes were mixed with LHCII to form LHCII-containing proteoliposomes and were purified via a sucrose gradient for a final homogeneous sample (Figure S1). The LHCII-to-lipid ratio was varied to produce LHCII proteoliposomes with an average number of LHCII, <N>, from less than one to ten. The ratios were selected based on lipid size, LHCII size, and the measured incorporation efficiency of LHCII^16^. The calculated <N> was confirmed through the advent of spectroscopic signatures of LHCII-LHCII interaction, as discussed below. Successful formation of the proteoliposomes was determined through dynamic light scattering (DLS), which showed a single peak indicating a monodisperse sample with a hydrodynamic radius of *∼*30-45 nm (Figure S2). The hydrodynamic radius of the proteoliposomes increased with LHCII content, likely due to the hydrophilic surface of LHCII (Table S1)^22^. Linear absorption and emission spectra of the purified LHCII proteoliposomes (Figures S3 and S5) showed the intact LHCII trimeric structure was maintained.

### LHCII quenching with increasing protein density

The fluorescence lifetime reports on the overall photophysical pathways, and several studies have established that it decreases in the presence of LHCII-LHCII interactions^14,15^. To characterize the overall dependence of the lifetime on the protein density, we performed ensemble time-resolved fluorescence measurements. The fluorescence decay traces were best fit with a bi-exponential function and the average lifetimes were calculated from these two terms (Figure S8). To investigate the distribution of fluorescence lifetime values, we performed single-molecule time-resolved fluorescence measurements (Figure S14-15). The lifetimes were extracted from the single-molecule data by fitting the fluorescence decay curves to single exponentials using maximum likelihood estimation (Figure 1a; SI Section 8). Although a bi-exponential function yielded the best fit for the ensemble decay curves, a single-exponential function was used owing to the lower signal-to-noise ratio in single-molecule data. Single-molecule measurements require high excitation fluences, and so light-harvesting proteins photodegrade during the observation time. To minimize the effect of photodegradation, the lifetime was only characterized immediately upon illumination, prior to any changes in the emission properties. The fluorescence lifetimes were measured and analyzed from many individual LHCII-proteoliposomes (*∼*100) for each sample to generate lifetime histograms as shown in Figure 1b.

**Figure 1.**
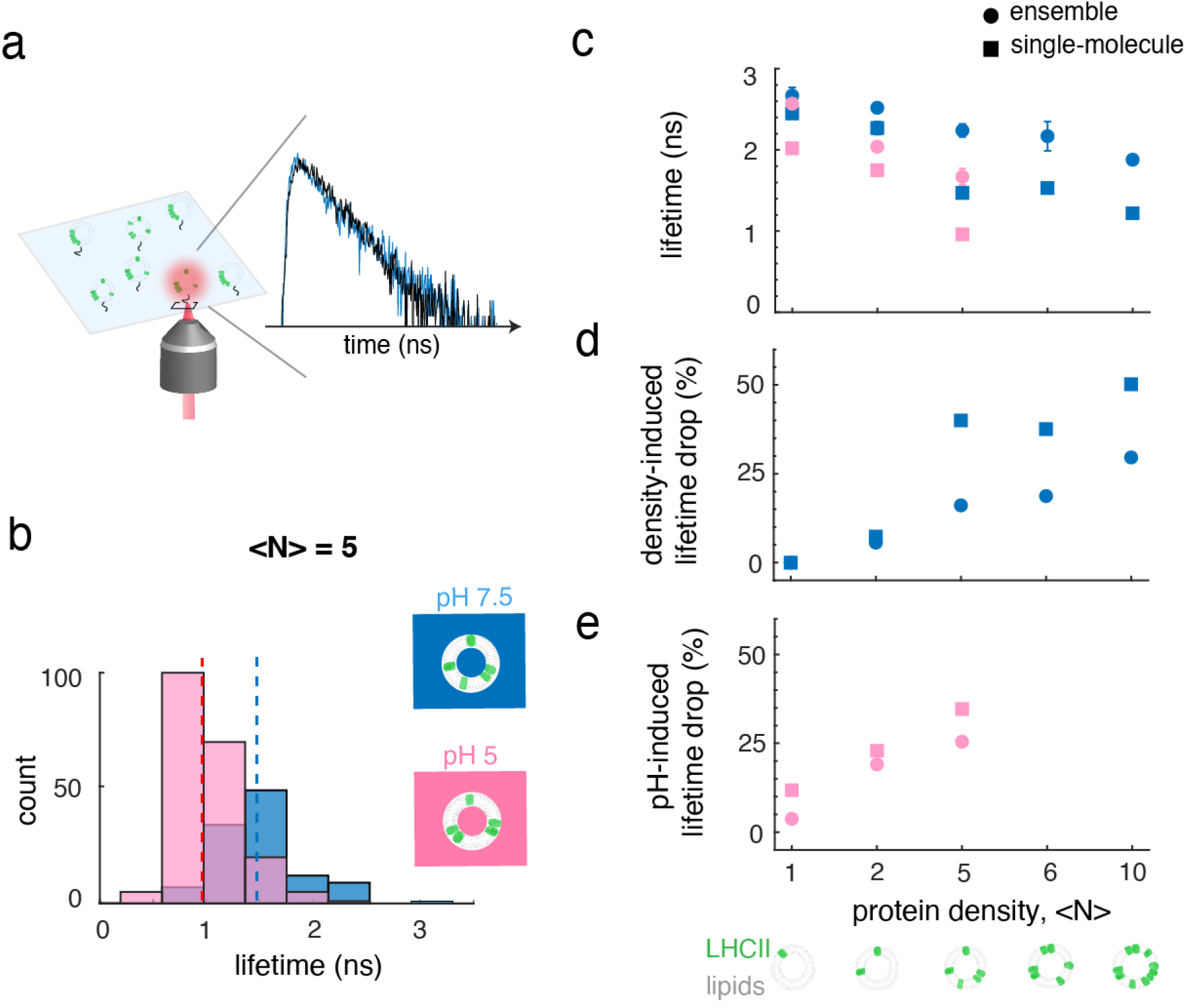
pH- and cluster-mediated quenching of LHCII in the liposome. (a) Schematic of LHCII-proteoliposomes immobilized on a coverslip for single-molecule measurements. A representative single-molecule fluorescence transient shows the fluorescence decay (blue) and its single-exponential fit (black). (b) Representative lifetime distributions obtained from single-molecule measurements of LHCII-proteoliposome samples with an average number of proteins per liposome (<N>) of 5 at pH 7.5 (blue) and 5.0 (pink). The dotted vertical lines are the medians of the distributions. (c) Lifetimes of the LHCII-proteoliposome samples at pH 7.5 (blue) and 5 (pink) at various protein densities. (d) Density- and (e) pH-induced lifetime drop of the proteoliposomes at different protein densities (see SI Sec 5). Circles and squares represent the values from the ensemble and single-molecule measurements, respectively.

The average (squares) and median (circles) lifetimes from the ensemble and single-molecule measurements, respectively, for all LHCII-proteoliposome samples at neutral and at acidic pH are shown in Figure 1c, Tables S2 and S3. The lifetime values decreased with protein density and with the pH drop in both ensemble and single-molecule measurements, revealing two types of quenching processes: (1) cluster-dependent quenching; and (2) pH-dependent quenching.

#### Cluster-dependent quenching

The ensemble measurements yielded an average lifetime of 2.67 ns for <N>= 1, consistent with previous work^23^. The single-molecule measurements yielded a slightly shorter median lifetime of 2.45 ns for <N>= 1, likely due to a small remaining effect of photodegradation. The lifetimes gradually decreased with protein density down to 1.88 ns for <N> = 10 for the ensemble data and 1.22 ns for <N> = 10 for the single-molecule data, all at pH 7.5. These values give an overall *∼*30% and *∼*50% reduction for the ensemble and single-molecule data, respectively (Figure 1d), corresponding a *∼*3-5% reduction per LHCII. Additionally, a *∼*20% difference between the ensemble and single-molecule measurements emerged at higher protein densities. Photodegradation typically results in the generation of quenchers, which can also quench neighboring proteins. The effect of photodegradation is thus expected to increase with protein density, consistent with these observations^23,24^.

#### pH-dependent quenching

The lifetimes were also measured after overnight incubation at pH 5 and decreased from the pH 7.5 values for the samples with <N>=1, 2 and 5 (Figure 1c,e). For LHCII-proteoliposomes with high (<N> >5) protein density, long incubation times (>150 hrs) are required for LHCII to fully equilibrate into clustered configurations in the membrane (SI Sec. 6). However, proteoliposomes become unstable after a few days, particularly at low pH, so the pH-dependence of these samples could not be investigated. The lifetime of LHCII-proteoliposome samples with <N>=1 decreased to an average of 2.57 ns for the ensemble data and a median of 2.02 ns for the single-molecule data, corresponding to a <10% reduction. This minor quenching at low pH could be either due to a pH-dependent conformational change of LHCII or a change in the local electrostatic environment.

Similar to the pH 7.5 data, the lifetimes at pH 5 gradually decreased as the protein density increased down to 1.67 ns for <N>=5 for the ensemble data and 0.96 ns for <N>=5 for the single-molecule data. The magnitude of the pH-induced lifetime reduction increased with protein density as shown in Figure 1e. The pH-induced reduction rose to *∼*25% and *∼*30% for the ensemble and single-molecule data, respectively (Figure 1e), for <N>=5, corresponding to a *∼*5% reduction per LHCII. The difference in magnitude between the ensemble and single-molecule data is likely due to the presence of photodegradation in single-molecule data, which is more significant at low pH. Overall, the larger reduction in lifetime with the pH drop for higher protein densities likely arises from the ability of these samples to adopt more clustered configurations.

### Extraction of LHCII-LHCII interaction energies

The quenching with increasing protein density arises from protein-protein interactions between LHCII. Thus, the fluorescence lifetime provides a reporter for LHCII-LHCII interactions, where shorter lifetimes indicate more (stronger) LHCII-LHCII interactions and longer lifetimes indicate fewer (weaker) LHCII-LHCII interactions. Based on this dependence, we developed an analysis method to quantitatively extract LHCII-LHCII interaction energies from the single-molecule lifetime distributions of the LHCII-proteoliposomes as illustrated in Figure 2 (SI Sec. 15).

**Figure 2.**
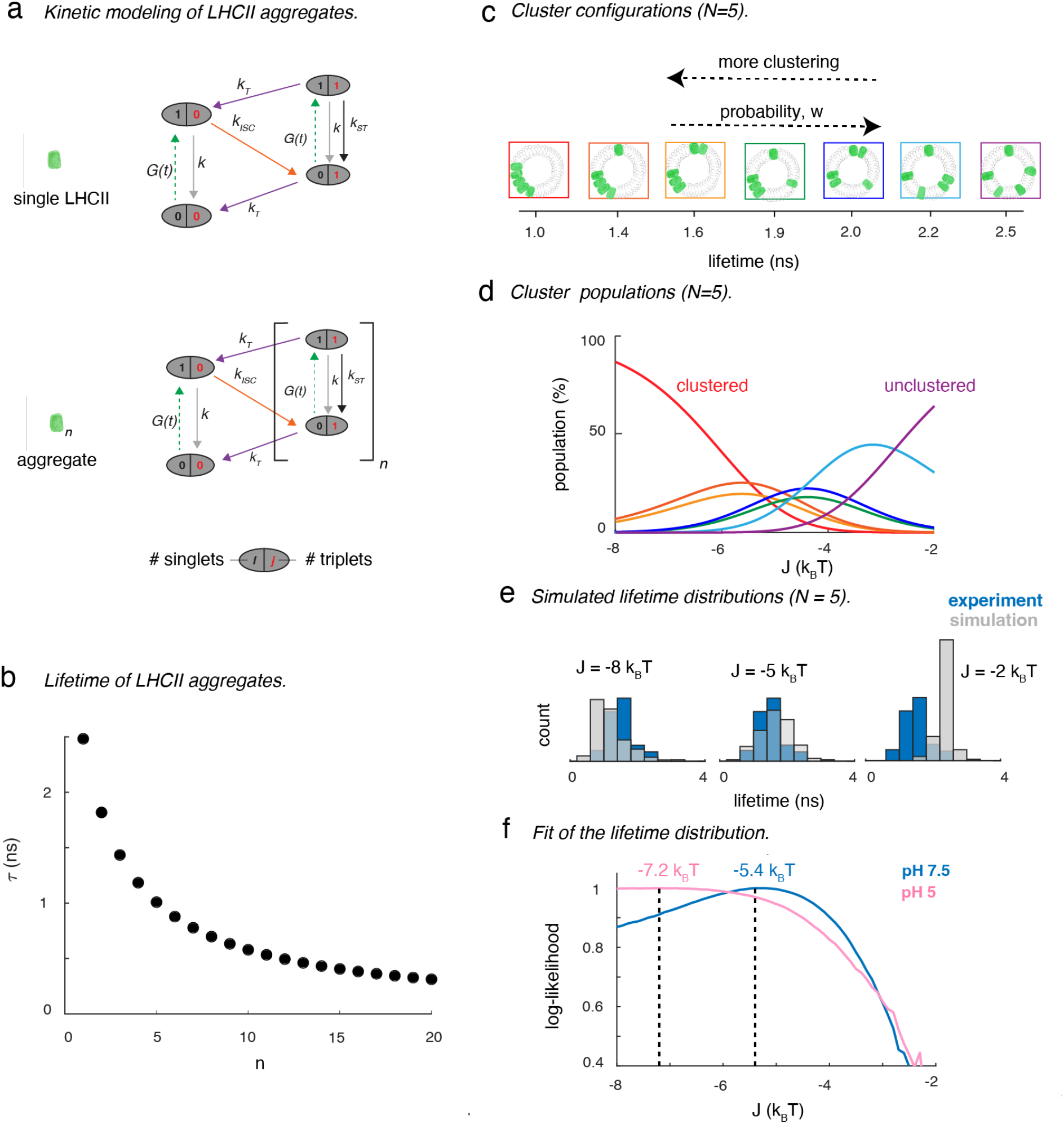
Extraction of LHCII-LHCII interaction energies from single-molecule lifetime data. (a) Stochastic kinetic model of the excited state relaxation pathways of isolated and clustered LHCII. The ovals represent states with different numbers of singlets (black) and triplets (red). *G*(*t*) is the excitation rate; *k* and *k*_*T*_ are rate constants for linear de-excitation from singlet and triplet states, respectively; *k*_*ISC*_ is the inter-system crossing rate-constant; and *k*_*ST*_ is the rate-constant for singlet-triplet annihilation. (b) Fluorescence lifetime computed from the model in (a) as a function of the number of LHCII complexes in a cluster, n. (c) All possible cluster configurations of LHCII-proteoliposomes upon the incorporation of a total number of proteins, N, of five and the lifetime of each configuration estimated from the model. (d) The populations of these seven configurations (color-matched to the boxes of the respective configurations in (c)) are displayed as a function of LHCII-LHCII interaction energy, *J*. (e) The relative populations of these configurations are utilized to obtain lifetime distributions (gray histograms) at *J* = -8, -5, and -2 *k*_*B*_*T*. The simulated distributions are overlaid with the experimental lifetime distribution (blue histograms). (f) Results of global fits of the simulated distributions to the experimental ones to extract *J* at pH 7.5 (blue) and 5 (pink).

In the first step of our method (Figure 2a), we developed a stochastic model adapted from Gruber et al.^25^ to predict the fluorescence lifetimes of LHCII clusters of different sizes. This model takes known photophysical properties of LHCII and incorporates the influence of singlet-triplet annihilation and the dependency of such annihilation process on cluster size (*n*), which arises from the ability of a triplet excitation to quench singlets in the surrounding LHCII within the cluster. Figure 2b displays the lifetime as a function of *n*. The model predicts a lifetime of 2.5 ns for *n* = 1, consistent with the experimental value of 2.7 ns for the <N>=1 proteoliposomes, which decreases to <0.5 ns for *n* = 20.

In the second step of our method (Figure 2c), we characterized different configurations of the same number of proteins (*N*) containing different cluster sizes (*n*). For a given number of proteins, the possible number of configurations (*m*) was computed. For instance, with N=5, seven different configurations are possible ranging from completely clustered to completely unclustered as illustrated in Figure 2c. The probability of formation (*w*) of each configuration was also computed based on their geometry, which depends on the number of proteins, the size of the proteins, and the size of the proteoliposome (SI Sec 12). As expected, *w* decreased for configurations with more clustering. The lifetime of each configuration was also calculated using the lifetime of its constituent clusters as described above.

In the third step, we quantified the equilibrium populations of each configuration, which depend both on the enthalpy and the entropy of the system. The enthalpy of the LHCII proteoliposome system can be mathematically described as a function of pairwise LHCII-LHCII interaction energy (*J*) while the entropy is related to the probability of formation (*w*). Figure 2d displays the population of different configurations for N=5 as a function of *J*.

In the fourth step, we generated simulated lifetime distributions from the number of proteins (*N*), the relative populations of the different configurations with a given *N*, and the lifetimes of each configuration (Figure S29 and S30). Figure 2e displays the simulated lifetime distributions (gray) overlaid with the experimental lifetime distribution (blue) for <N> = 5 at *J* of -8, -5 and -2 *k*_*B*_*T*. The simulated lifetime distribution is in good agreement with the experiment at *J* = -5 *k*_*B*_*T*, indicating the value of *J* is in that vicinity.

In the final step, the simulated lifetime distributions for all <N> were globally fit to the experimental ones at each pH using maximum likelihood estimation (MLE) to extract *J* (SI Sec 15). Figure 2f shows the log-likelihood estimates as a function of *J* (fits for individual samples are shown in Figure S31). The best fits for the LHCII-LHCII interaction energy were obtained at -5.4 and -7.2 *k*_*B*_*T* for the samples at pH 7.5 and pH 5, respectively. The negative values signify that LHCII-LHCII pairwise interaction energies are attractive in nature, and the attraction is strengthened by *∼*2 *k*_*B*_*T* upon the pH drop.

### Free-energy driving force for LHCII clustering in liposomes

Using the identified interaction energies, we quantified the free energy driving force associated with the reorganization of LHCII into clusters upon a pH drop. We derived the following expression for the free energy change (Δ*G*) for the transition of LHCII-proteoliposomes from neutral to low pH:

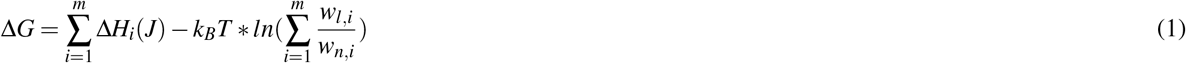

where Δ*H*_*i*_(*J*) is the change in enthalpy associated with *i*-th configurations and *w*_*l,i*_ and *w*_*n,i*_ are the probabilities of such configurations at pH 5.0 and 7.5, respectively. The first term of the right-hand side of the above equation represents the enthalpy change and the second term represents the change in entropy for this transition. For <N>=5, enthalpy and entropy changes were found to be -12.7 *k*_*B*_*T* and -4.5 *k*_*B*_, respectively. These values show that this transition is mildly exothermic in nature and associated with a reduction in entropy presumably due to a more restricted organization with clusters. From the changes in enthalpy and entropy, the Gibbs free energy change is quantified as -8.2 *k*_*B*_*T* for <N>=5. For other protein densities, the free energy changes ranged from -2 to -19 *k*_*B*_*T* (Figure 3a, Table S9). While the magnitudes of the changes in entropy, enthalpy and Gibbs free energy all increased with protein density, the change in entropy only increased slightly whereas the change in enthalpy increased significantly, and so the free energy became even more dominated by the enthalpic contribution at high protein density.

**Figure 3.**
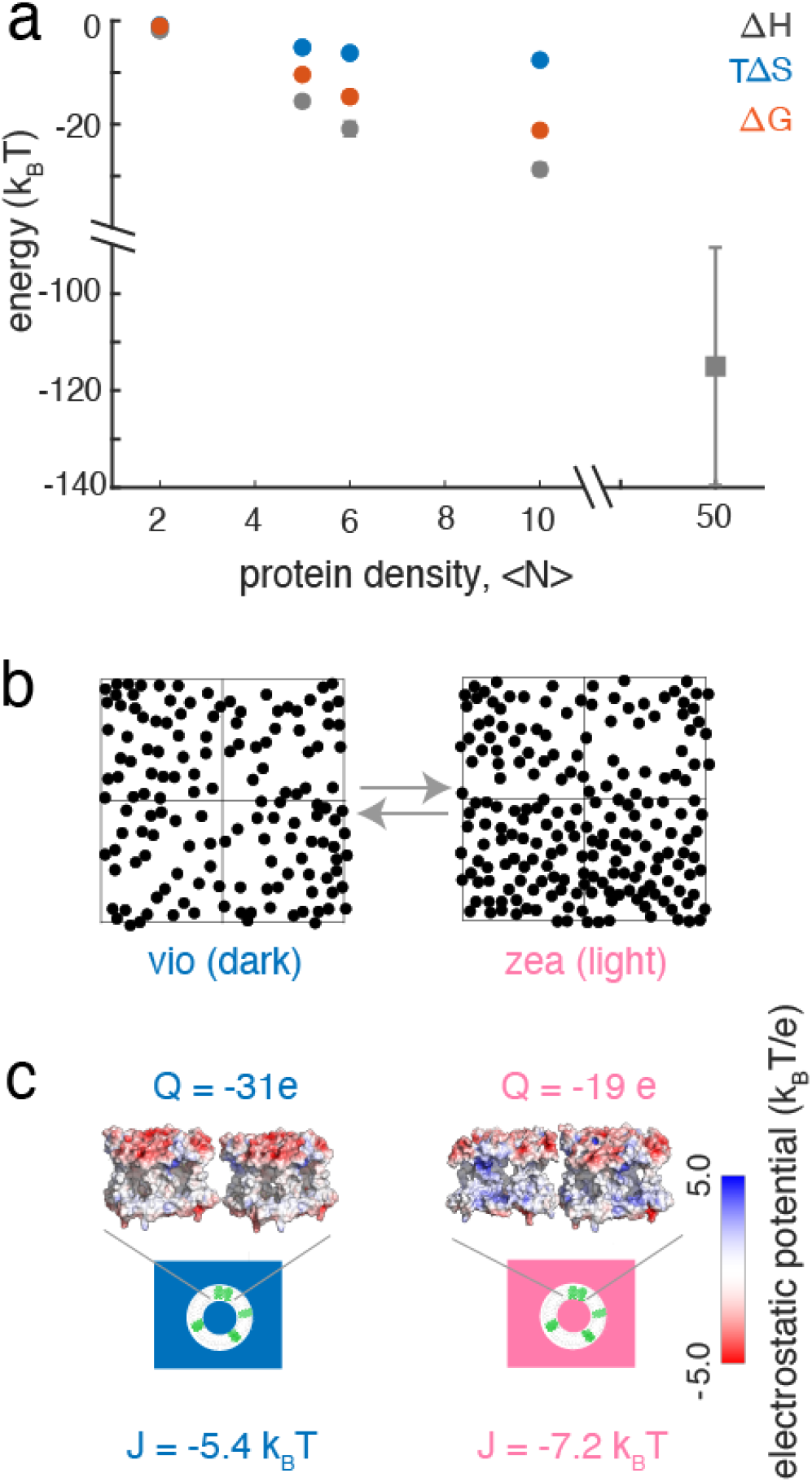
Free-energy change of pH-induced clustering of LHCII. (a) The changes in enthalpy (ΔH), entropy (ΔS) and Gibbs free-energy (ΔG) in the LHCII proteoliposomes (circles) and chloroplasts (square) for the pH-induced or high light-induced clustering. The error bars are the standard errors. (b) Masks produced by thresholding the freeze-fracture electron micrographs of the intact spinach chloroplasts of the dark-adapted (vio, dark) and light-treated (zea, light) samples reported in^20^. The square boxes are the *∼* 88 nm *×* 88 nm regions over which the thermodynamics parameters involving a transition from dark to light organizations were calculated. (c) Schematic of lateral interactions between two LHCII apo-proteins at pH of 7.5 (left) and 5 (right). LHCII proteins are shown as potential energy surfaces generated by the Adaptive Poisson-Boltzmann solver in Pymol with the net surface charges of the proteins (Q) also displayed.

### Free-energy driving force for enhanced LHCII clustering in the intact chloroplast

To quantify the free-energy change that drives the enhancement of clustering under high light in vivo, we analyzed the freeze-fracture electron micrographs of the intact spinach chloroplasts reported by Johnson et al.^20^ (SI Sec. 18). The dark-adapted (vio, dark) and light-adapted (zea, light) membranes were assigned to light-harvesting and dissipative states, respectively, based on the reported NPQ values^20^. Particle picker transforms of the electron micrographs were used to identify the LHCII organization in the membrane (Figure 3b, SI Sec. 18). The transforms were analyzed for each *∼* 88 nm *×* 88 nm region of the micrographs, which is similar in surface area to the proteoliposomes. Based on the extracted membrane organizations and the LHCII interaction energies extracted from the LHCII-proteoliposome experiments, we calculated ΔH associated with the transition from the light-harvesting to a dissipative state. The average value of ΔH in the chloroplast scaled similarly to the values from the LHCII-proteoliposomes. For instance, in proteoliposome with <N> = 10, we calculated ΔH *∼* -28 *k*_*B*_*T* associated with the transition from high to low pH. Now, with a five-fold increase in the density (<N> = 50) in the chloroplast, the change in enthalpy is found to be *∼* -115 *k*_*B*_*T*. The large error associated with the ΔH value in the chloroplast reflects the stochasticity in protein organization in the different regions of the membrane.

## DISCUSSION

### Molecular-level origin of LHCII-LHCII interactions

We quantified the LHCII-LHCII interaction energy as approximately -5 *k*_*B*_*T* at neutral pH and at least -7 *k*_*B*_*T* at acidic pH (2f, SI Sec 15, Fig S31). The negative sign implicates these interaction energies are indeed attractive. Schneider et al. estimated LHCII-LHCII stacking interactions across the stromal gap as -4 *k*_*B*_*T* ^26^ based on the electrostatic attraction between two correspondingly charged parallel plates in an aqueous solution of counter ions. Therefore, the lateral LHCII-LHCII interaction obtained in the current work is *∼*30% stronger than the predicted stacking interaction, consistent with the formation of protein networks laterally within the thylakoid membrane^20,27,28^.

The net interactions between the proteins can be viewed as a balance between electrostatic repulsion and van der Waals attraction forces^21,29,30^. The net charge of free LHCII trimers is -31e, which arises from the carboxyl group associated with the glutamic acid and aspartic acid residue side chains^21,29^. The van der Waals attraction arises due to the interactions between uncharged molecules with permanent and/or instantaneously induced electrostatic dipole moment. As the LHCII-LHCII interaction energy is attractive in nature, the attractive van der Waals force must overpower the repulsive electrostatic force. Similarly, a recent computational study showed PSII-LHCII interactions are attractive due to the same balance of van der Waals and electrostatic forces^31^. Because the majority of charges in LHCII are localized at the terminals, the transmembrane part of the protein is largely hydrophobic in nature. This charge distribution minimizes the electrostatic repulsion between LHCII trimers, thereby leading to enhancedvan der Waals attraction. The net charge and therefore overall surface charge density is highly dependent on the membrane lipid composition, local ionic strength, and charged residues, and so the specific amplitudes of the two opposing forces are likely adjusted by these factors in vivo. For example, phosphorylation of LHCII introduces additional negative charges, which likely increase the electrostatic repulsion among LHCII and weaken the LHCII-LHCII lateral interactions^21,32,33^. Alternatively, with an abundance of cations in the medium, the effective net charge of LHCII is decreased and the magnitude of the repulsive interactions is also decreased, strengthening the LHCII-LHCII interactions. Indeed, the effect of screening on LHCII interactions has been demonstrated by Kirchhoff and coworkers using positively charged ions (K^+^, Mg^2+^)^21^.

What could be the molecular-level origin of enhanced interaction at low pH? van der Waals forces depend on the size of the proteins, inter-protein distances and the dielectric constant of the medium, all of which are nearly constant between low and neutral pH. We propose that the net electrostatic repulsion between the proteins decreases at low pH due to the screening effect of H^+^, the protonation of negatively charged lipids in the membrane, or the protonation of certain negatively charged amino acid residues of LHCII exposed to the solvent. Our estimation of the net surface charge of LHCII proteins (*Q*) in low and neutral pH supports this hypothesis (Figure 3c, Sec SI 15). The surface charge of LHCII is mostly located at its edges facing the stromal and lumenal sides of the membrane while the central transmembrane region is largely neutral. *Q* was found to decrease from -31e at pH 7 to -19e at pH 5, corresponding to a 38% reduction of negative charge at low pH consistent with the measured *∼*30% increase in pairwise interaction energy.

### Enhanced LHCII clustering at low pH

Intense work in the last few decades has found that LHCII is a major site for qE in higher plants and that ΔpH is necessary for its trigger^12,20,34–36^. Numerous previous studies have also revealed the tendency of LHCII proteins to cluster^14,15,17,19^. Our statistical thermodynamical modeling explains the clustering of LHCII and its enhancement at low pH, providing a basis for the organization of LHCII and its dependence on qE conditions. LHCII-LHCII interactions are required for cluster formation, as the population of clustered configurations in the proteoliposomes is near zero when the interaction energy is not present, *i*.*e*., at *J*=0 (Figure 2d). At neutral pH where moderate interactions are present (*J* = -5.4 *k*_*B*_*T*), a variety of configurations co-exist primarily containing a mixture of clustered and unclustered LHCII. The interaction energies produce an enthalpic driving force (ΔH) that overcomes the entropic driving force (ΔS) associated with isolated LHCII to induce moderate clustering. At low pH where stronger interactions are present (*J* = -7.2 *k*_*B*_*T*), the population of clustered configurations significantly increases due to the exponential dependence on enthalpic energy (*H*(*J*), see Methods, Eqn. 5). For instance, in the LHCII-proteoliposomes with <N>=5, the percentage of the completely clustered configuration, where all five LHCII form a single cluster, is 25% at the neutral pH and increases to 74% at low pH (Figure S33). Ultimately, the difference in interaction energies leads to an enthalpic driving force (ΔH) that induces a transition from moderately clustered configurations at neutral pH to heavily clustered configurations at low pH.

### LHCII clustering from a thermodynamic perspective

We quantified the changes in enthalpy (ΔH), entropy (ΔS), and free energy (ΔG) for the enhanced LHCII clustering associated with the transition from a light-harvesting (neutral pH) to dissipative (low pH) state (Figure 3a). The change in enthalpy is favorable (Δ*H* < 0) owing to the attractive LHCII-LHCII interactions. In contrast, the change in entropy is unfavorable (Δ*S* <0), as expected from the reduced probabilities of highly clustered configurations. The larger magnitude of the enthalpic term as compared to the entropic term leads to a Δ*G* < 0, enabling spontaneous or thermodynamically feasible enhancement of LHCII clustering at low pH (Table S9). These results reveal that the enhanced clustering of LHCII in the liposome or natural thylakoid membrane at low pH is primarily driven by enthalpy.

The magnitude of the enthalpic term increases dramatically with protein density owing to the additional possible interactions, which scale combinatorially with the number of LHCII. In contrast, the magnitude of the entropic term only changes minimally owing to the relatively dilute LHCII density (Figure 3a). That is, the similar number of configurations available for all the proteoliposome samples in this low-density limit leads to similar entropic values. The increase in the magnitude of ΔH, and thus the magnitude of ΔG, makes the clustering process more exergonic in nature as protein density increases. However, at the even higher protein densities found in the chloroplast (equivalent to <N>=50), the magnitude of the free energy change may be different compared to its liposome counterpart. The thylakoid environment is extremely crowded with different proteins, and so the entropic term for LHCII clustering is likely smaller owing to the smaller effective area. The clustering in vivo is likely more complicated owing to the presence of other factors, such as the carotenoid zeaxanthin, ions, charged lipids, and interactions with other proteins such as PsbS. Indeed, dimeric PsbS is thought to monomerize at low pH^37^, increasing the entropy of the overall system. Therefore, ΔS may also induce other membrane reorganizations during NPQ similar to its role in state transitions^38^. The complexity of the native photosynthetic membrane may alter the interactions, organization, and thus thermodynamics associated with NPQ, changing the specific values of the thermodynamic parameters. While the values may be modified, the free energy driving forces quantified here are on the order of a few *k*_*B*_*T* s, and thus represent values large enough to induce a membrane reorganization under qE conditions but low enough so that their effect can be reversible through thermal motion and likely regulated by external factors.

## CONCLUSION

In this article, by employing single-molecule measurements of LHCII-proteoliposome samples at various protein densities and modeling the photophysics of LHCII-clusters, we quantified the lateral LHCII-LHCII interaction energies at neutral and acidic pH. The interaction energy at neutral pH is found to be approximately -5 *k*_*B*_*T*, which creates a strong but reversible binding to form clusters of various sizes. The interaction energy is *∼*30% stronger at pH 5 compared to at pH 7.5, which drives a transition from moderate clustering to a strong clustering state. Thermodynamic analysis established that the change in free energy associated with the enhanced clustering at low pH is largely enthalpy-driven and on the order of 10s *k*_*B*_*T* s, which is strong enough to form clusters but low enough to be reversible and regulated by external factors. These values capture the underlying mechanisms behind the long established yet previously unexplained reorganization of the plant membrane, which is thought to play an active and important role in the regulation of photosynthesis. Overall, this work presents a framework to analyze protein clusters using equilibrium statistical thermodynamics that can be extended to investigate the organization of other membrane proteins.

## Methods

### Preparation of LHCII-proteoliposomes

LHCII proteins used in this work were purified from the dark-adapted spinach as described in details elsewhere^23^. The detailed protocol for the preparation of LHCII proteoliposome at neutral pH (pH, 7.5) and their characterizations are reported in the SI (SI Sec. 3) of this article and also in^23^. For the low pH measurements, proteoliposome prepared at neutral pH was incubated for *∼* 10-12 hrs. at (20 mM MES, 40 mM NaCl, pH 5). The size of the incubated liposomes was monitored at the different phases of incubation using dynamic light scattering (Figure S10).

### Ensemble Measurements

Absorption and emission spectra of the LHCII samples were collected using an Epoch Microplate Spectrophotometer (BioTek) and a Cary Eclipse Fluorescence Spectrophotometer, respectively (Figure S3-S5). The samples were syringe filtered (GE Healthcare Life Sciences, pore size 0.22 *μ*m) to discard any aggregates before the measurements.

The fluorescence decays were measured with a TCSPC module (Time Tagger 20, Swabian Instruments). For excitation, a tunable fiber laser (FemtoFiber pro, Toptica Photonics, 80 MHz repetition rate, 130 fs pulse duration, 610 nm, 4 nm full-width half maximum (fwhm)), passed through a pinhole and directed into a home-built confocal microscope. The excitation was focused by an oil-immersion objective (UPLSAPO100XO, Olympus, NA 1.4) onto the sample placed on a coverslip. The emission of the sample was collected through the same objective and separated from the excitation using a dichroic (ZT647rdc, Chroma) and bandpass filter (ET700/75m, Chroma). The fluorescence decay was fit by iterative re-convolution with a bi-exponential function using the measured instrument response function (IRF) of the system with a home-built MATLAB code. The IRF was measured by the scattered signal to be *∼* 400 ps (fwhm). The average lifetime values were obtained through an intensity-weighted average of the fitted bi-exponential lifetime constants (Table S2, Figure S8).

### Single-molecule Measurements

The filtered samples were diluted to *∼* 15 pM LHCII in 20 mM HEPES, 40 mM NaCl, pH 7.5 (for neutral buffer) or 20 mM MES, 40 mM NaCl, pH 5 (for acidic buffer). For the measurement at neutral pH (pH 7.5), we used an enzymatic scavenging mixture containing 2.5 mM protocatechuic acid (PCA) and 25 nM protocatechuate-3,4-dioxygenase (PCD). A mixture of 20 nM Pyranose Oxidase from Microorganism (Creative Enzymes, Shirley, NY), 1.2 *μ*g/ml of catalase (Sigma-Aldrich) and 100 *μ*M glucose was used as scavenging solution for low pH measurements. For the low pH measurements, argon gas was also used to avoid intense photobleaching. The proteoliposomes were immobilized on a homemade or commercial (Bio 01 low-density glass coverslip, 22 *×* 22 mm, Microsurfaces Inc.) biotinylated coverslip via neutravidin-biotin interactions. The method of single-molecule data collection and analysis has been described in the SI Sec. 7-8.

### Modeling

#### Stochastic modeling of LHCII clusters

We developed a model of the excited-states of LHCII clusters to predict the fluorescence lifetimes. Upon photoexcitation of their singlet states, chlorophyll molecules relax via several competing pathways^39^: (1) radiative decay from the Chl singlet states (1/*k ∼* 5 ns); (2) inter-system crossing to Chl triplet states (1/*k*_*ISC*_ *∼* 10 ns), which subsequently transfer to Cars. Under the high excitation fluence and repetition rate of single-molecule measurements, there can be a significant accumulation of such Car triplets. The Car triplets de-excite any singlet excited states via singlet-triplet annihilation^40^, which has been extensively described in LHCII^25,41^. In this study, the average number of proteins in the proteoliposome is ten or below, and these discrete numbers of trimers are best described with a stochastic model, similar to Gruber et al.^25^ (SI Sec. 13). The contribution of singlet-singlet annihilation is insignificant (<2.7%) under the excitation fluences used in this work and therefore was excluded from the model.

Owing to the influence of S-T annihilation, the excited-state lifetime of a single LHCII depends on the triplet concentration. Assuming a random distribution of triplet quenchers over LHCII aggregates and the S-T annihilation process being independent of their spatial coordinates, the dependence of lifetime on the triplet concentration ([*T*]) can be calculated^40,41^:

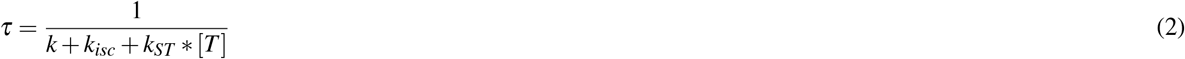

where *k, k*_*isc*_ and *k*_*ST*_ are the rate constants of the radiative decay, inter-system crossing and singlet-triplet annihilation, respectively^25^. The equation also assumes that the triplet population has reached a quasi-stationary condition as the decay of triplet states (microseconds) is much slower than the singlet state decay (nanoseconds). The rate constants used in this model were obtained from^25^ and shown in the SI Sec 13. At higher illumination intensities, the triplet state population ([*T*]) increases, resulting in an enhanced quenching of the singlets (Eqn.2). In the first step of the analysis, we described the excited-state lifetime of LHCII and LHCII clusters (Figure 2a). To validate our model, we performed intensity-dependent single-molecule measurements of unclustered LHCII (Figure S27). The lifetime decreased with illumination intensity, as expected. The model predictions matched reasonably well with the median values of the experimental lifetime distributions (Table S7).

The amount of S-T annihilation is also dependent on the cluster size^42^. As the size of the cluster increases, the singlets can be quenched by triplets from other LHCII, increasing the probability of the S-T annihilation. In the small clusters investigated here, we assume a triplet present in a single LHCII can quench the excitations in the entire cluster. In this limit, the total number of effective triplets or magnitude of triplet quenching can be modeled to increase linearly with the cluster size. Therefore, for a cluster consisting of *n* number of LHCII molecules, the lifetime of its singlet excited state is given as:

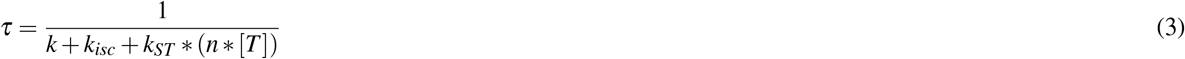

where, [*T*] is the triplet concentration in an isolated LHCII. Next, we used this model to calculate the excited-state lifetime as a function of the size of the LHCII cluster (*n*) as displayed in Figure 2b. Stochastic modeling of LHCII photophysics, characterization of the different clusters and their equilibrium populations, generation of simulated lifetime distributions at different protein-protein interaction energies (*J*), and extraction of *J* based on maximum likelihood estimation have been described in detail in the Supplementary Information section of this article.

#### Cluster configurations and populations

At high protein densities, LHCII clusters form on liposomes that exhibit quenching of the fluorescence emission^15,42^. The cluster formation mimics the in vivo formation of LHCII arrays under high light conditions^20,43^. In the second step of the analysis (Figure 2c), we characterized the clusters by employing a classical statistical mechanics description of an interacting system^44^. The configurations for a given number of proteins, *N*, can be obtained by solving the equation below (SI Sec 10)^44^:

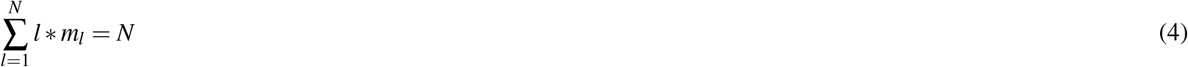

where the system forms *m*_*l*_ number of clusters of *l* proteins. The *m*-matrix constructed from the solutions to the equation above contains the possible configurations of the *N* proteins, *i*.*e*., the sizes of the clusters. For example, when *N* = 5, seven different configurations of the five LHCII are possible as shown in Figure 2c. In the most clustered configuration (left), all the LHCII form one cluster of five proteins whereas in the least clustered configuration (right), the proteins are all separated. As the number of proteins increases, the number of configurations also increases (Figure S16b). For instance, when *N* = 10, 42 different configurations are possible.

However, the probability (*w*) of forming these different types of configurations varies depending on the total number of proteins involved (*N*), the geometry of clusters, i.e. the connectivity of the proteins in the configurations, and also the available area to form the clusters. For instance, the probability of forming a cluster of two proteins in a liposome can be estimated as follows: assuming one LHCII protein is already on the liposome, if we incorporate another protein, what is the chance that the second protein will be within 1 nm distance (which is our definition of the cluster) to the first one to form a cluster of two proteins vs. both remains unclustered. It turns out to be that assuming zero interactions between the proteins, the probability of forming two clusters on a liposome with a radius of 25 nm is only 0.006. The probability of forming different types of configurations on a spherical surface (*w*) is discussed in detail in the Supplementary Information section of this article (SI Sec 12).

The lifetime of each configuration is computed by first calculating the lifetime for each cluster within the configuration using the model values displayed in Figure 2c and second combining the values to describe the full configuration. Figure 2b shows that the configurations with more clustering are characterized by a shorter lifetime and lower probability of formation (*w*).

At equilibrium, the complex interplay between entropy and enthalpy determines the population of a configuration^45^. In a clustered system like the LHCII-proteoliposomes, the enthalpy (*H*) can be mathematically represented as a function of the pairwise interaction energy between proteins. We introduce this parameter, the pairwise LHCII-LHCII interaction energy, as *J*. For a given number of LHCII in the liposome, the equilibrium population of the *i*^*th*^ configuration (*p*_*i*_) depends on the probability of forming that configuration (*w*_*i*_) and its total enthalpy (*H*_*i*_(*J*)) in the following way:

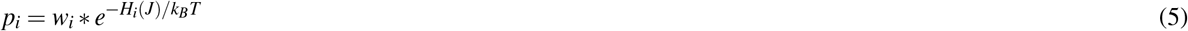

where, *k*_*B*_ and *T* are the Boltzmann constant and temperature, respectively.

The enthalpy (*H*(*J*)) of a configuration depends on how the LHCII proteins are clustered in that configuration. For instance, in a configuration of five LHCII where two and three proteins are clustered (Figure 2c), *H*(*J*)is given as:

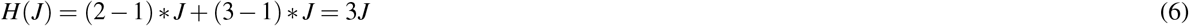

Figure 2d displays the population of the seven configurations for N=5 as a function of *J*. When *J* is close to zero, mostly unclustered configurations are favored as there is no energetic driving force to form clusters. In contrast, when *J* is large (close to -10 *k*_*B*_*T*), clustered configurations are favored. At an intermediate *J*, populations with both clustered and unclustered configurations coexist.

#### Simulated lifetime distribution and extraction of LHCII-LHCII interaction energy

For a liposome with a certain number of LHCII embedded in it, we simulate the possible cluster configurations, their excited state lifetimes, and relative populations from our model. This enables us to generate the lifetime distribution of that sample (SI Sec 15). First, the number of proteins incorporated into a given liposome, *N*, is selected with a probability given by a Poissonian distribution with the average value, <N>, for the sample. All possible configurations for *N* are modeled. For instance, if an *N* of five is selected, we model all seven configurations as shown in Figure 2c. The relative population of these configurations depends on the pairwise LHCII-LHCII interaction energy (*J*) as discussed above (Figure 2d). Next, at a certain *J*, a possible configuration is selected and weighted by its relative population. The lifetime of that configuration is computed from the stochastic model as discussed above (also in SI Sec 13). This whole process is repeated multiple times to generate a lifetime distribution for that sample at a particular *J* (Figure 2e).

### Free-energy change calculation

At equilibrium, the Gibbs free energy (Δ*G*) change for a transition from neutral to low pH is described in the following way:

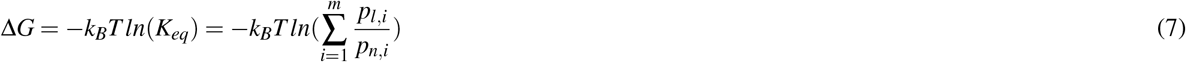

where, *K*_*eq*_ is the equilibrium constant, *p*_*n,i*_ and *p*_*l,i*_ are the population of the *i*-th configurations at the neutral (pH 7.5) and low pH (pH 5), respectively. *m* is the number of possible configurations for a given number of proteins embedded in the liposome as discussed above. Now, the population of the *i*^*th*^ configuration, *p*_*i*_ is given as shown in Eqn. 5. Inserting Eqn. 5 in Eqn. 7 and slight re-arranging we obtain Eqn. 1 which is used to compute the free-energy change of the transition from neutral to low pH (see also SI Sec. 17).

## Acknowledgements

This work is supported by the Human Frontiers Science Program (Award #RGY0076 to MPJ and GSSC). PM, MH, and GSSC also acknowledge support from the U.S. Department of Energy, Office of Science, Office of Basic Energy Sciences, Division of Chemical Sciences, Geosciences, and Biosciences under Award # DE-SC0018097 to GSSC. GSSC thanks the CIFAR Bio-inspired Solar Energy program for support.

## Author contributions

PM and GSSC conceived the project. PM, MH and TD performed the experiments. PM and MH analyzed the data. PM and GSSC wrote the original draft of the paper. GSSC and MPJ supervised the work. All authors reviewed the manuscript.

## References

1. Dill, K. A., Bromberg, S. & Stigter, D. Molecular driving forces: statistical thermodynamics in biology, chemistry, physics, and nanoscience (Garland Science, 2010).

2. Garcia, H. G., Kondev, J., Orme, N., Theriot, J. A. & Phillips, R. Thermodynamics of biological processes. In Methods in enzymology, vol. 492, 27–59 (Elsevier, 2011).

3. Stevers, L. M., de Vink, P. J., Ottmann, C., Huskens, J. & Brunsveld, L. A thermodynamic model for multivalency in 14-3-3 protein–protein interactions. J. Am. Chem. Soc. 140, 14498–14510 (2018).

4. Sartori, P. & Leibler, S. Lessons from equilibrium statistical physics regarding the assembly of protein complexes. Proc. Natl. Acad. Sci. 117, 114–120 (2020).

5. Chong, S.-H. & Ham, S. Dynamics of hydration water plays a key role in determining the binding thermodynamics of protein complexes. Sci. Reports 7, 8744 (2017).

6. van Oort, B., van Hoek, A., Ruban, A. V. & van Amerongen, H. Equilibrium between quenched and nonquenched conformations of the major plant light-harvesting complex studied with high-pressure time-resolved fluorescence. The J. Phys. Chem. B 111, 7631–7637 (2007).

7. Santabarbara, S., Horton, P. & Ruban, A. V. Comparison of the thermodynamic landscapes of unfolding and formation of the energy dissipative state in the isolated light harvesting complex II. Biophys. journal 97, 1188–1197 (2009).

8. Bassi, R. & Dall’Osto, L. Dissipation of light energy absorbed in excess: the molecular mechanisms. Annu. Rev. Plant Biol. 72, 47–76 (2021).

9. Ruban, A. V. Nonphotochemical chlorophyll fluorescence quenching: Mechanism and effectiveness in protecting plants from photodamage. Plant physiology 170, 1903–1916 (2016).

10. Giovagnetti, V. & Ruban, A. V. The evolution of the photoprotective antenna proteins in oxygenic photosynthetic eukaryotes. Biochem. Soc. Transactions 46, 1263–1277 (2018).

11. Muller, P., Li, X.-P. & Niyogi, K. K. Non-photochemical quenching. a response to excess light energy. Plant physiology 125, 1558–1566 (2001).

12. Nicol, L., Nawrocki, W. J. & Croce, R. Disentangling the sites of non-photochemical quenching in vascular plants. Nat. plants 5, 1177–1183 (2019).

13. Manna, P. & Schlau-Cohen, G. S. Photoprotective conformational dynamics of photosynthetic light-harvesting proteins. Biochimica et Biophys. Acta (BBA)-Bioenergetics 1863, 148543 (2022).

14. Natali, A. et al. Light-harvesting complexes (LHCs) cluster spontaneously in membrane environment leading to shortening of their excited state lifetimes. J. Biol. Chem. 291, 16730–16739 (2016).

15. Tutkus, M. et al. Aggregation-related quenching of LHCII fluorescence in liposomes revealed by single-molecule spectroscopy. J. Photochem. Photobiol. B: Biol. 218, 112174 (2021).

16. Tutkus, M. et al. Fluorescence microscopy of single liposomes with incorporated pigment–proteins. Langmuir 34, 14410–14418, DOI: 10.1021/acs.langmuir.8b02307 (2018).

17. Akhtar, P., Görföl, F., Garab, G. & Lambrev, P. H. Dependence of chlorophyll fluorescence quenching on the lipid-to-protein ratio in reconstituted light-harvesting complex II membranes containing lipid labels. Chem. Phys. 522, 242–248 (2019).

18. Shukla, M. K. et al. A novel method produces native light-harvesting complex II aggregates from the photosynthetic membrane revealing their role in nonphotochemical quenching. J. Biol. Chem. 295, 17816–17826, DOI: https://doi.org/10.1074/jbc.RA120.016181 (2020).

19. Wilson, S., Li, D.-H. & Ruban, A. V. The structural and spectral features of light-harvesting complex II proteoliposomes mimic those of native thylakoid membranes. The J. Phys. Chem. Lett. 13, 5683–5691, DOI: 10.1021/acs.jpclett.2c01019 (2022). Doi: 10.1021/acs.jpclett.2c01019.

20. Johnson, M. P. et al. Photoprotective energy dissipation involves the reorganization of photosystem II light-harvesting complexes in the grana membranes of spinach chloroplasts. The Plant Cell 23, 1468–1479 (2011).

21. Puthiyaveetil, S., Van Oort, B. & Kirchhoff, H. Surface charge dynamics in photosynthetic membranes and the structural consequences. Nat. Plants 3, 1–9 (2017).

22. Liu, Z. et al. Crystal structure of spinach major light-harvesting complex at 2.72 Å resolution. Nature 428, 287–292 (2004).

23. Manna, P., Davies, T., Hoffmann, M., Johnson, M. P. & Schlau-Cohen, G. S. Membrane-dependent heterogeneity of LHCII characterized using single-molecule spectroscopy. Biophys. J. 120, 3091–3102 (2021).

24. Wang, Q. & Moerner, W. Dissecting pigment architecture of individual photosynthetic antenna complexes in solution. Proc. Natl. Acad. Sci. 112, 13880–13885 (2015).

25. Gruber, J. M., Chmeliov, J., Krüger, T. P., Valkunas, L. & Van Grondelle, R. Singlet–triplet annihilation in single LHCII complexes. Phys. Chem. Chem. Phys. 17, 19844–19853 (2015).

26. Schneider, A. R. & Geissler, P. L. Coexistence of fluid and crystalline phases of proteins in photosynthetic membranes. Biophys. journal 105, 1161–1170 (2013).

27. Su, X. et al. Structure and assembly mechanism of plant C_2_S_2_M_2_-type PSII-LHCII supercomplex. Science 357, 815–820 (2017).

28. Shen, L. et al. Structure of a C_2_S_2_M_2_N_2_-type PSII–LHCII supercomplex from the green alga Chlamydomonas reinhardtii. Proc. Natl. Acad. Sci. 116, 21246–21255 (2019).

29. Barber, J. Influence of surface charges on thylakoid structure and function. Annu. Rev. Plant Physiol. 33, 261–295 (1982).

30. Chow, W. S., Kim, E.-H., Horton, P. & Anderson, J. M. Granal stacking of thylakoid membranes in higher plant chloroplasts: the physicochemical forces at work and the functional consequences that ensue. Photochem. & Photobiol. Sci. 4, 1081–1090 (2005).

31. Mao, R., Zhang, H., Bie, L., Liu, L.-N. & Gao, J. Million-atom molecular dynamics simulations reveal the interfacial interactions and assembly of plant PSII-LHCII supercomplex. RSC advances 13, 6699–6712 (2023).

32. Bellafiore, S., Barneche, F., Peltier, G. & Rochaix, J.-D. State transitions and light adaptation require chloroplast thylakoid protein kinase STN7. Nature 433, 892–895 (2005).

33. Wood, W. H. & Johnson, M. P. Modeling the role of LHCII-LHCII, PSII-LHCII, and PSI-LHCII interactions in state transitions. Biophys. J. 119, 287–299 (2020).

34. Saccon, F., Giovagnetti, V., Shukla, M. K. & Ruban, A. V. Rapid regulation of photosynthetic light harvesting in the absence of minor antenna and reaction centre complexes. J. experimental botany 71, 3626–3637 (2020).

35. Ruban, A. V. & Wilson, S. The mechanism of non-photochemical quenching in plants: localization and driving forces. Plant Cell Physiol. 62, 1063–1072 (2021).

36. Nicol, L. & Croce, R. The PsbS protein and low pH are necessary and sufficient to induce quenching in the light-harvesting complex of plants LHCII. Sci. Reports 11, 7415 (2021).

37. Fan, M. et al. Crystal structures of the PsbS protein essential for photoprotection in plants. Nat. structural & molecular biology 22, 729–735 (2015).

38. Jia, H., Liggins, J. R. & Chow, W. S. Entropy and biological systems: Experimentally-investigated entropy-driven stacking of plant photosynthetic membranes. Sci. Reports 4, 1–7 (2014).

39. Ostroumov, E. E., Khan, Y. R., Scholes, G. D. et al. Photophysics of photosynthetic pigment-protein complexes. In Non-Photochemical Quenching and Energy Dissipation in Plants, Algae and Cyanobacteria, 97–128 (Springer, 2014).

40. Valkunas, L., Trinkunas, G., Liuolia, V. & Van Grondelle, R. Nonlinear annihilation of excitations in photosynthetic systems. Biophys. journal 69, 1117–1129 (1995).

41. Zaushitsyn, Y., Jespersen, K. G., Valkunas, L., Sundström, V. & Yartsev, A. Ultrafast dynamics of singlet-singlet and singlet-triplet exciton annihilation in poly (3-2’-methoxy-5’octylphenyl) thiophene films. Phys. Rev. B 75, 195201 (2007).

42. Barzda, V. et al. Fluorescence lifetime heterogeneity in aggregates of LHCII revealed by time-resolved microscopy. Biophys. journal 81, 538–546 (2001).

43. Lambrev, P. H. et al. Importance of trimer–trimer interactions for the native state of the plant light-harvesting complex II. Biochimica et Biophys. Acta (BBA)-Bioenergetics 1767, 847–853 (2007).

44. Sinha, S. K. Classical statistical mechanics of interacting system. In Introduction to statistical mechanics, 204–239 (Alpha Science Int’l Ltd., 2005).

45. Schneider, A. R. & Geissler, P. L. Coarse-grained computer simulation of dynamics in thylakoid membranes: methods and opportunities. Front. Plant Sci. 4, 555 (2014).

