## Supplemental Information for "Energetic driving force for LHCII clustering in plant membranes"

**4 Trypsin Digest** **11**

**5 Lifetime Measurements at Low pH** **12**

**6 Kinetics of Cluster Formation in Liposomes** **14**

**7 Single-molecule Measurements** **16**

**8 Single-molecule Data Analysis** **18**

**9 Density- and pH-induced lifetime drop in LHCII proteoliposome.** **18**

**10 Clustering of interacting proteins.** **19**

**11 Determination of the equilibrium population of cluster configurations from simulation.** **20**

**12 Analytical solution of the probability of cluster formation on liposome.** **21**

**13 Photokinetic modeling of cluster-mediated lifetime quenching of LHCII in lipid environment.** **26**

**14 Intensity dependence of excited state lifetime in the isolated LHCII complexes** **31**

**15 Extraction of LHCII-LHCII interaction energy from lifetime data.** **32**

**16 Surface Charge Calculations** **37**

**17 Free energy change in pH-driven clustering in LHCII proteoliposome.** **38**

**18 Free energy change in light-driven clustering in the chloroplast.** **40**

### 1 Purification of LHCII

LHCII trimers used in this study were purified from spinach. The details of the LHCII purification and the characterizations are provided in [9]. Briefly, fresh market spinach leaves were dark-adapted at 4°C overnight then blended in ice-cold grinding medium (330 mM Sorbitol, 10 mM $Na_4P_2O_7$, 5 mM $MgCl_2$, 2 mM D(+) iso-ascorbate, pH 6.5). The homogenate was filtered once through two layers of muslin and one layer of cotton wool. The filtrate was centrifuged at 4000 x g for 20 minutes and the pellet resuspended in wash medium (330 mM Sorbitol, 10 mM MES, pH 6.5) followed by centrifugation at 4000 x g for 20 minutes. The pellet was resuspended in resuspension medium (330 mM Sorbitol, 40 mM MES, 5 mM $MgCl_2$, pH 6.5). Break medium (5 mM $MgCl_2$, 40 mM MES, pH 7.6) was added to triple the original volume to lyse any unbroken chloroplasts. The osmotic potential was restored after 45 seconds with the addition of an equal volume of osmoticum medium (660 mM Sorbitol, 40 mM MES, 5 mM $MgCl_2$, pH 6.5). The sample was then centrifuged at 4000 x g for 20 minutes and the thylakoid membranes were then resuspended in a small volume of storage medium (300 mM Sucrose, 10 mM HEPES, 5 mM $MgCl_2$, pH 7.5). Thylakoids were solubilized at a concentration of 1 mg/ml chlorophyll in 0.55% n-Dodecyl- β-D-Maltopyranoside (β-DDM) in the dark on ice for 10 minutes, with occasional mixing. Partially solubilized thylakoids were centrifuged at 1000 x g for 10 minutes to pellet any insoluble material. The supernatant was removed and centrifuged 38,000 x g for 40 minutes to pellet Photosystem II membrane fragments (known as BBYs). BBYs with a final chlorophyll concentration of 0.5 mg/ml were solubilized in 10mM HEPES (pH 7.5), 0.5% n-Hexadecyl-β-D-Maltopyranoside (HDM), 0.1 % β-DDM for 30 minutes at room temperature, with occasional mixing. The solubilized BBYs were centrifuged at 22,000 x g for 10 minutes and the supernatant was retained. LHCII was purified from solubilized BBYs via sucrose density gradient centrifugation. Solubilized BBYs were loaded onto a continuous 650 mM freeze/thaw sucrose gradient in a buffer of 20 mM HEPES, pH 7.5, 0.06% (w/v) glyco-diosgenin (GDN) in SW32 Ti rotor tubes (Beckman Coulter, Inc.). Fractionation of protein complexes was performed by centrifugation at 175,000 x g in an SW32 Ti swinging bucket rotor (Beckman Coulter, Inc.) for 28-32 hours at 4°C. After centrifugation, trimeric LHCII was harvested using a peristaltic pump. Finally, LHCII samples were flash-frozen in liquid nitrogen and stored at -70°C.

### 2 Preparation of LHCII-proteoliposomes

For the preparation of liposomes, a thylakoid lipid mixture containing 50% monogalactosyldiacylglycerol (MGDG), 30% digalactosyldiacylglycerol (DGDG), 10% sodium salt of L-α-phosphatidylglycerol (Soy PG) and 10% sulfoquinovosyldiacylglycerol (SQDG) was dissolved in 1 ml of 7:3 chloroform: MeOH. Biotinylated lipids (sodium salt of 1,2-dioleoyl-sn-glycero-3-phosphoethanolamine-N-(cap biotinyl), 18:1 Biotinyl Cap PE, Avanti Polar Lipids, Inc.) were added to the mixture for a final ratio of 1:250 (biotinylated lipid: thylakoid lipid). The lipids were completely dried down using $N_2$ gas and transferred to a vacuum desiccator for 2-3 hrs. The lipid films prepared this way are stored at -70°C and used subsequently as needed. The dried lipid mix was resuspended in a working buffer (20 mM HEPES, 40 mM NaCl, pH 7.5) by vortexing vigorously. The lipid mixture was subjected to 8 freeze/thaw cycles to break down large multilamellar vesicles. The vesicles were then extruded through a polycarbonate membrane with 50 nm (diameter) pore ∼21-times using an extrusion kit (Avanti Polar Lipids, Inc.) to generate unilamellar vesicles with uniform sizes. 0.03% β-DDM was gently mixed with the sample to destabilize the unilamellar liposomes and the mixture was left for 30 minutes with occasional gentle mixing. The desired amount of LHCII was added to the sample containing empty liposomes to obtain a lipid:protein molar ratio of 7000, 5000, 2500, 1000, 880, and 510 with the average number of proteins per liposome ($\langle N \rangle$) $<1$, 1, 2, 5, 6 and 10, respectively.

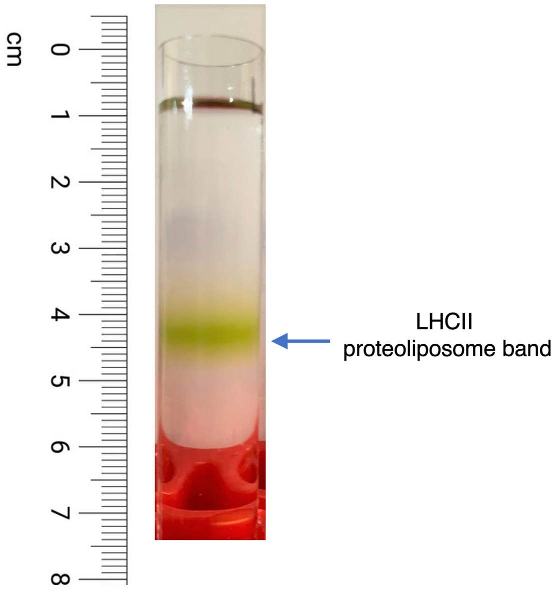

Figure S1: A representative sucrose gradient showing a LHCII proteoliposome band for <N>=5.

This number is based on a $\sim$ 20-25% incorporation of LHCII in proteoliposome as shown here [20]. The reaction mixture was left for 1 hr. in the dark with occasional gentle mixing. Bio-beads (Bio-beads SM-2 adsorbent media, Bio-Rad Laboratories, Inc.) were added in a step-wise manner into the solution for 2-3 hrs. to extract detergent molecules from the solution.

A 5-step sucrose gradient (15%, 20%, 25%, 30%, and 35%) was prepared in SW41 ultracentrifuge tubes (Beckman Coulter, Inc.) using a peristaltic pump. LHCII-proteoliposomes were loaded onto the sucrose gradients which were then centrifuged at 150,000 x g for 14 hrs. to remove liposome-free LHCII aggregates. After centrifugation, the green band corresponding to LHCII-proteoliposomes was collected from the gradient using a peristaltic pump and subjected to further ensemble characterization and single-molecule measurements within one week.

### 3 Characterization of LHCII proteoliposomes

#### 3.1 Dynamic light scattering

For DLS measurements, the samples were filtered to remove large aggregates. DynaPro Nanostar (Wyatt Technologies) was used for DLS measurements with illumination at 658 nm. A commercial software (Dynamics 7.1.9.3) was used for the data analysis in DLS to report the intensity distribution of the scattered particles. The diameter of the proteoliposome samples are displayed in Table S1.

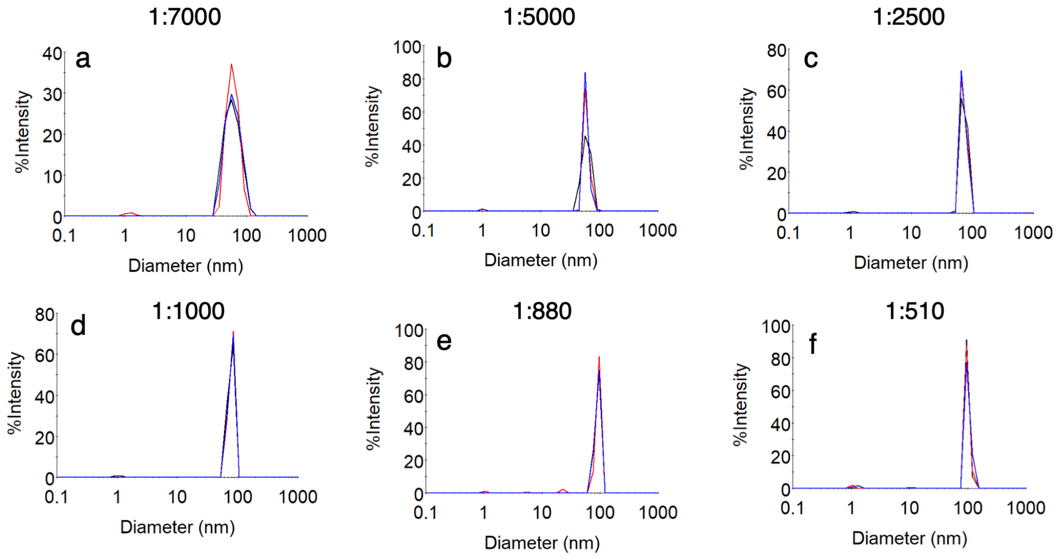

Figure S2: Size distribution of LHCII-proteoliposomes with various protein-to-lipid ratios obtained from dynamic light scattering (DLS) measurements. Blue, red and black lines represent the technical replicates of the measurements.

Table S1: Diameter of LHCII proteoliposomes with various protein-to-lipid ratios measured by dynamic light scattering. The numbers in the parenthesis are the standard deviations from independent measurements (n = 3).

| protein:lipid | <N> | diameter (nm) |
|---|---|---|
| 1:7000 | <1 | 61 (*1*) |
| 1:5000 | 1 | 60 (*0.9*) |
| 1:2500 | 2 | 73 (*0.9*) |
| 1:1000 | 5 | 79 (*0.7*) |
| 1:880 | 6 | 96 (*4*) |
| 1:510 | 10 | 99 (*2*) |

#### 3.2 Absorption and emission

Absorption and emission spectra of the LHCII samples were collected using an Epoch Microplate Spectrophotometer (BioTek) and a Cary Eclipse Fluorescence Spectrophotometer, respectively. The samples were syringe filtered (GE Healthcare Life Sciences, pore size 0.22 $\mu$m) to discard any aggregates before the measurements.

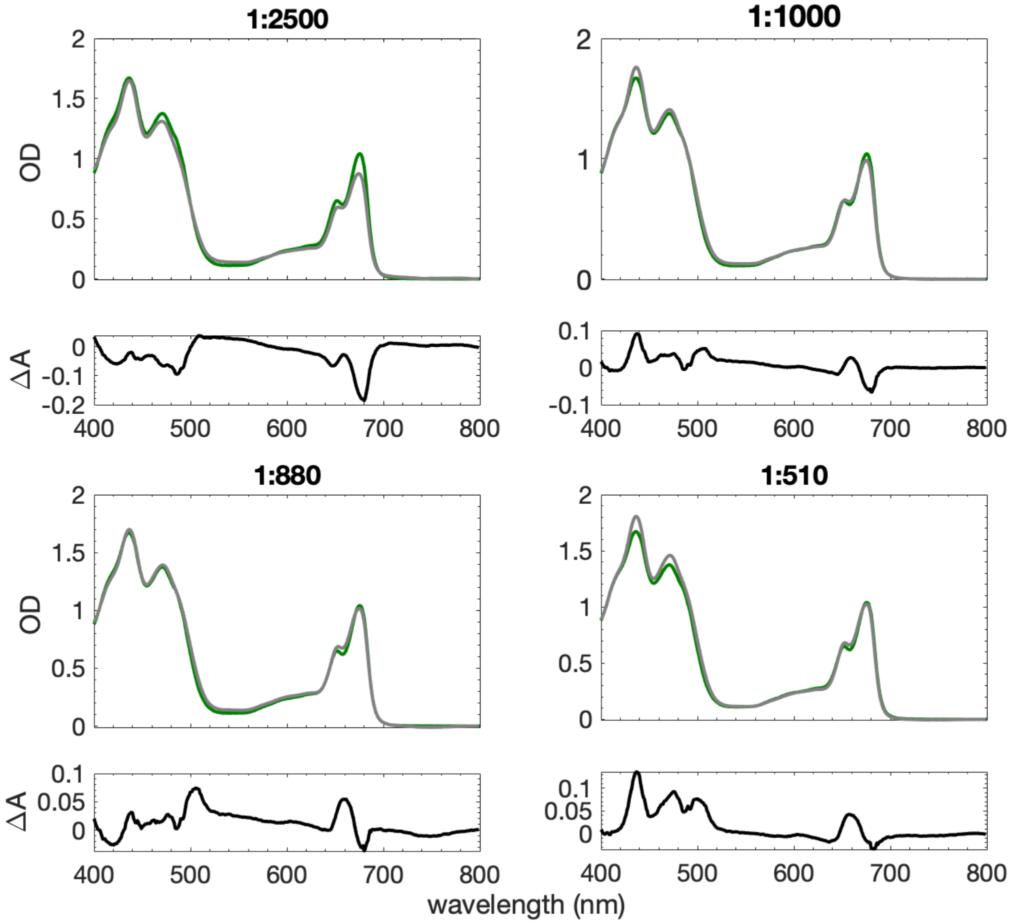

Figure S3: Absorption spectra of LHCII in detergent (10mM HEPES, 20 mM NaCl, 0.06% (w/v) GDN, pH 7.5) and incorporated in liposomes with protein-to-lipid ratio of 2500, 1000, 880 and 510. The difference spectra ($\Delta A = OD_{liposome} - OD_{detergent}$) are also displayed below each of the corresponding absorption spectra. Absorption spectra are normalized at 405 nm.

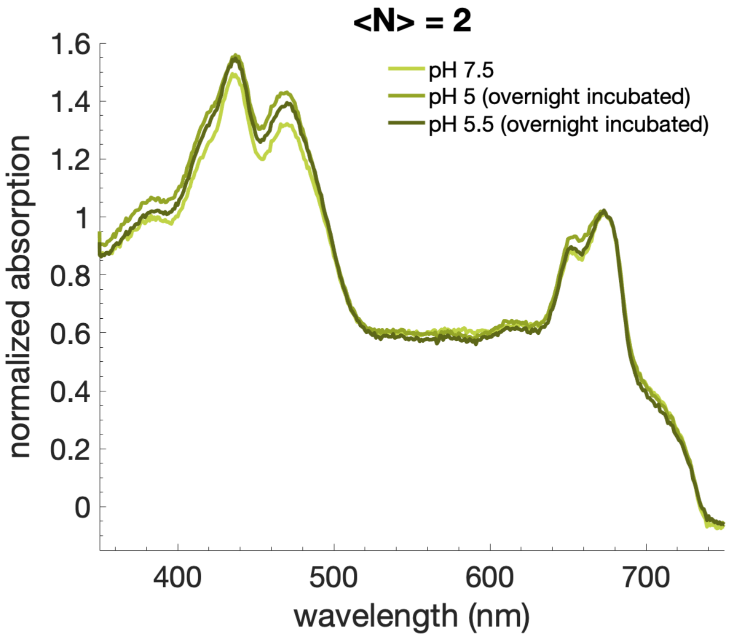

Figure S4: Absorption spectra of LHCII proteoliposome with <N>=2 incubated overnight at low pH.

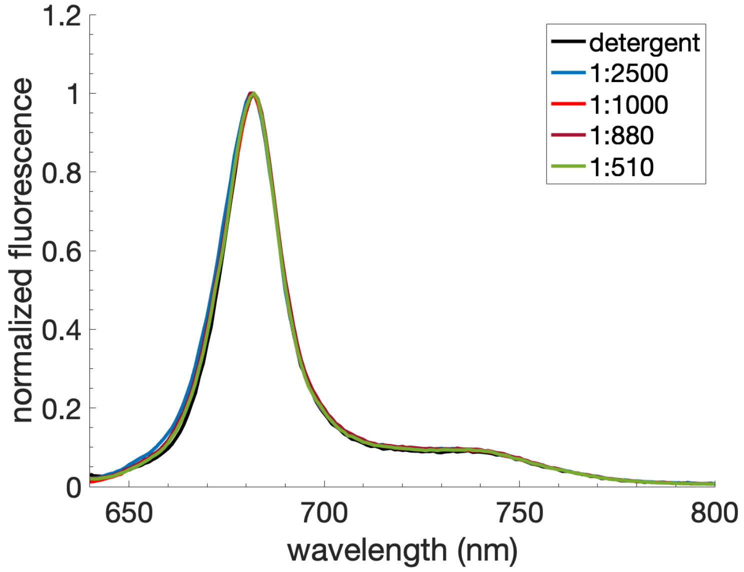

Figure S5: Normalized emission spectra of LHCII in detergent (10mM HEPES, 20 mM NaCl, 0.06% (w/v) GDN, pH 7.5) and incorporated in liposomes with protein-to-lipid ratios of 2500, 1000, 880 and 510.

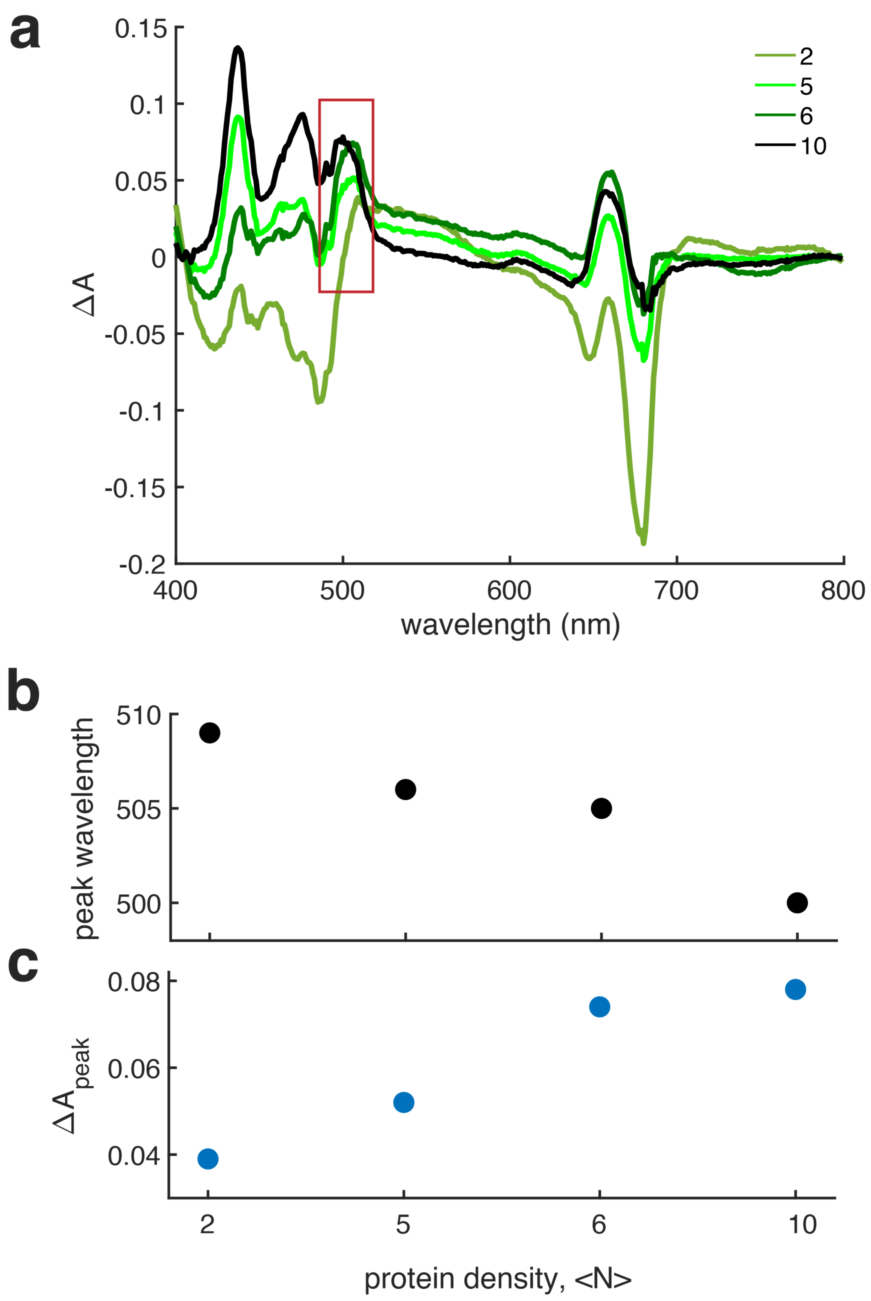

Figure S6: (a) Difference spectra ($\Delta A = OD_{liposome} - OD_{detergent}$) of the LHCII proteoliposome with varying protein density. The peak wavelengths around the 500 nm region of the difference spectra (within the box shown in (a)) and $\Delta A$ at that peak wavelengths are displayed in (b) and (c), respectively.

##### 3.3 Excited state lifetime

The fluorescence decays were measured with a TCSPC module (Time Tagger 20, Swabian Instruments). For excitation, a tunable fiber laser (FemtoFiber pro, Toptica Photonics, 80 MHz repetition rate, 130 fs pulse duration, 610 nm, 4 nm full-width half maximum (fwhm)), passed through a pinhole and directed into a home-built confocal microscope. The excitation was focused by an oil-immersion objective (UPLSAPO100XO, Olympus, NA 1.4) onto the sample placed on a coverslip. The emission of the sample was collected through the same objective and separated from the excitation using a dichroic (ZT647rdc, Chroma) and bandpass filters (ET700/75m, Chroma and ET690/120x, Chroma). The fluorescence decay was fit by iterative re-convolution with a bi-exponential function using the measured instrument response function (IRF) of the system with a home-built MATLAB code. The IRF was measured by the scattered signal to be $\sim$ 400 ps (fwhm). The average lifetime values were obtained through an intensity-weighted average of the fitted bi-exponential lifetime constants.

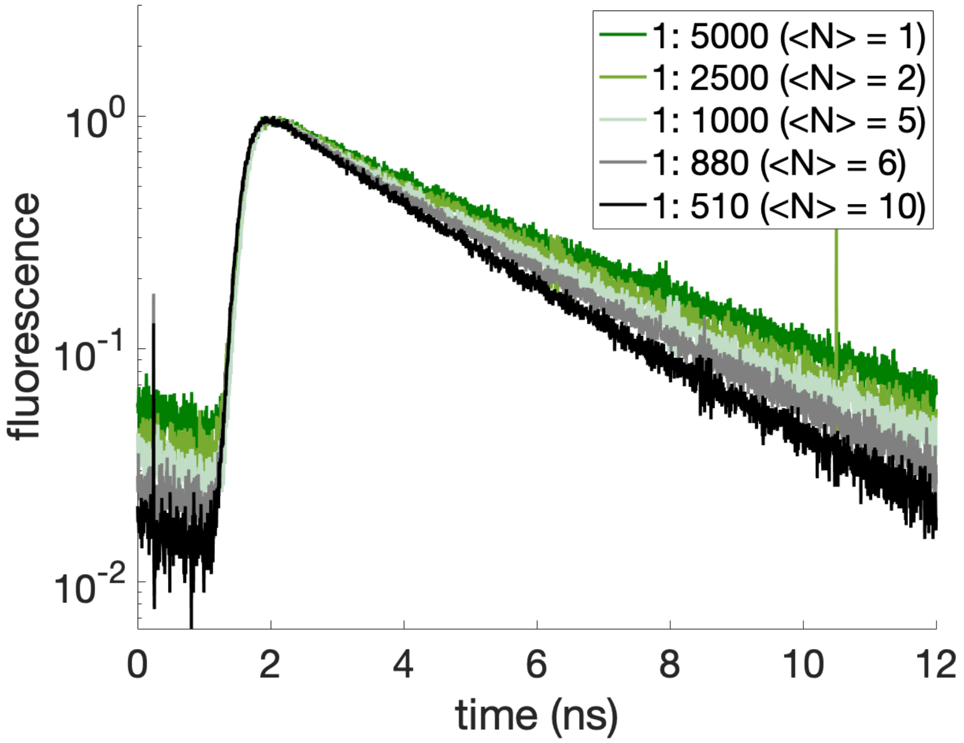

Figure S7: **Ensemble lifetime of LHCII proteoliposome samples**. Normalized fluorescence lifetime decay of LHCII embedded in liposomes with various protein-to-lipid ratios. The protein-to-lipid ratios used in this study are 1:5000, 1:2500, 1:1000, 1:880 and 1:510. The average number of LHCII (<N>) in liposomes in these samples are 1, 2, 5, 6 and 10 respectively. The fluorescence lifetime decay gets shortened upon increasing protein density in liposomes.

Table S2: Ensemble lifetime of LHCII in liposome at various protein densities (<N>). The values in the parenthesis are the errors obtained from standard deviations of 2-3 replicates.

| protein:lipid | <N> | pH | $a_1$ (%) | $\tau_1$ (ns) | $a_2$ (%) | $\tau_2$ (ns) | $\tau_{avg}$ (ns) |
|---|---|---|---|---|---|---|---|
| 1:7000 | <1 | | 87 | 3.00 | 13 | 0.61 | 2.69 |
| 1:5000 | 1 | | 81 | 3.24 | 19 | 0.24 | 2.67 (*0.10*) |
| 1:2500 | 2 | 7.5 | 81 | 3.00 | 19 | 0.37 | 2.52 (*0.01*) |
| 1:1000 | 5 | | 78 | 2.70 | 22 | 0.61 | 2.24 (*0.08*) |
| 1:880 | 6 | | 78 | 2.64 | 22 | 0.44 | 2.17 (*0.18*) |
| 1:510 | 10 | | 76 | 2.37 | 24 | 0.35 | 1.88 |
| 1:5000 | 1 | | 83 | 3.02 | 17 | 0.34 | 2.57 (*0.06*) |
| 1:2500 | 2 | 5.0 | 73 | 2.65 | 27 | 0.39 | 2.04 (*0.06*) |
| 1:1000 | 5 | | 67 | 2.24 | 33 | 0.47 | 1.67 (*0.10*) |

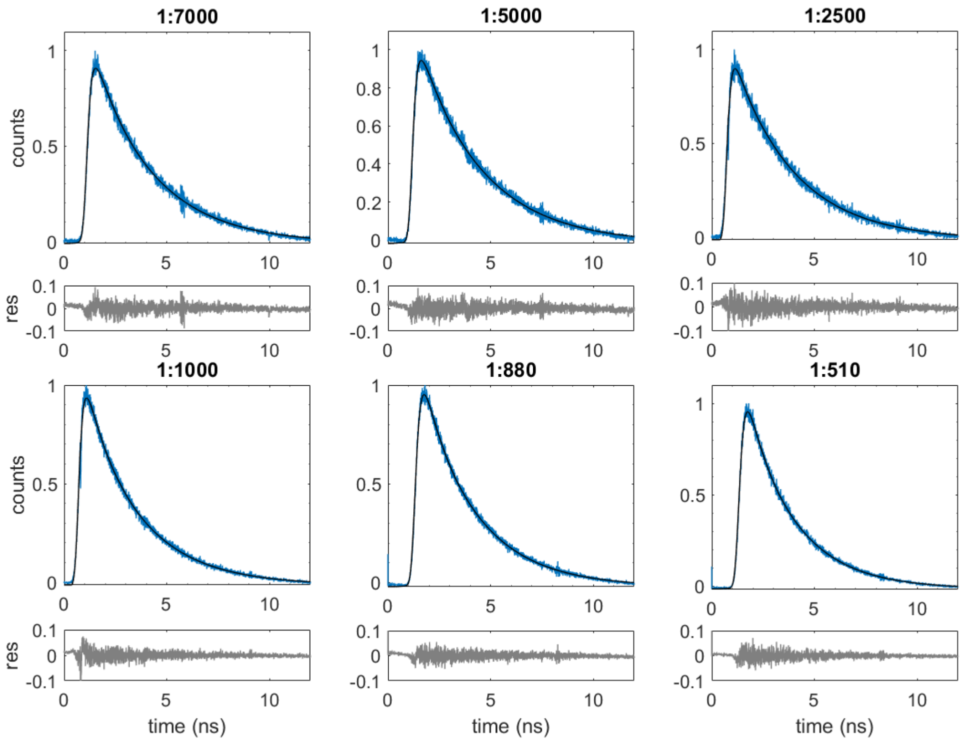

Figure S8: The ensemble lifetime decay of LHCII-proteoliposomes at different protein-to-lipid ratios. The figures display the fluorescence decay (blue lines) of the samples and their corresponding fits (black lines) along with the residuals. The decays are fitted with bi-exponential functions employing the non-linear least square-based method. The fit results are displayed in Table S2.

### 4 Trypsin Digest

Trypsin is an enzyme that cleaves arginine and lysine residues in a protein sample. In LHCII, trypsin selectively cleaves the N-terminus of the protein [2]. If LHCII inserts directionally into a liposome, then the LHCII should either be all uncleaved or totally cleaved after trypsin digestion. If it inserts randomly, we should see a mixture of cleavage products. Previous studies of LHCII in liposomes have reported mixed results [2, 12]. We performed trypsin digest experiments on the LHCII proteoliposomes used in this work to confirm orientation.

In brief, LHCII proteoliposome samples with $<N> = 5$ were mixed with trypsin in a 5:1 mass ratio and heated on a Thermomixer at 37°C for up to 2 hours. SDS-PAGE was used to visualize the reaction products. Standard protocols were used with the modification of not adding 2-mercaptoethanol prior to the heating step. 12% Mini-PROTEAN® TGX gels from Bio-Rad were found to have the best separation of the cleavage products.

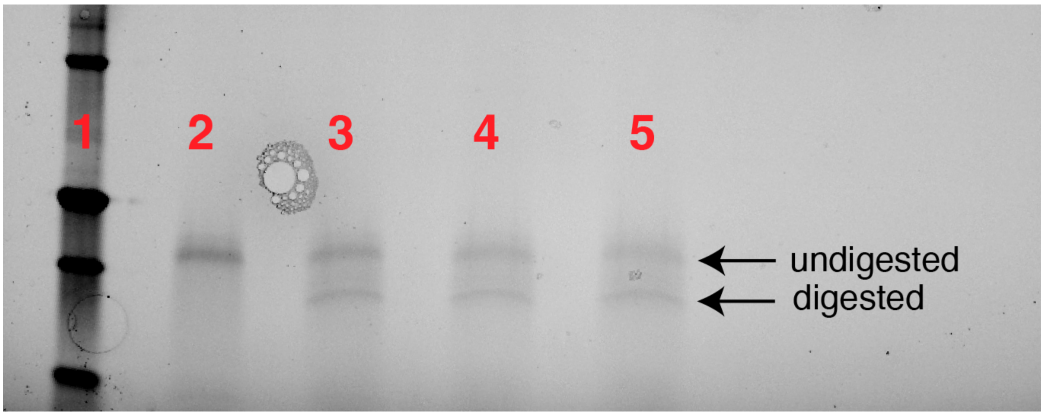

Figure S9: SDS-PAGE gel of trypsin digest products reveals random insertion of LHCII in liposome. The lanes are as follows: 1) Ladder. 2) LHCII proteoliposome pre-digestion. 3) Digestion products after 30 minutes. 4) Digestion products after 1 hour. 5) Digestion products after 2 hours.

### 5 Lifetime Measurements at Low pH

Protein incorporation efficiency in liposomes depends on the pH of the solvent and therefore liposomes prepared with the same protein-to-lipid ratio but at different pH differ in protein density. In order to disentangle the pH-induced and density-induced effects in the quenching of LHCII, it is paramount that the protein density in the liposome remains the same. Therefore, to overcome the uncertainty of protein density at neutral and low pH, we incubated the proteoliposome samples at pH 5 buffer rather than preparing the liposome at this low pH. We monitored the change in excited state lifetimes of the proteoliposomes over time to identify an equilibrium point (Figure S11). We also verified that the proteoliposome samples remain intact by performing dynamic light scattering (DLS) at different phases of the incubation process (Figure S10). It was found that the lifetime of LHCII-proteoliposome samples reached equilibrium after $\sim$ 10 hrs. of incubation and remained stable in the course of single-molecule data collection (S11 b,c).

As discussed in the main text, the enhanced quenching at low pH is driven by increased clustering of LHCII proteins which require diffusion of the proteins on the liposome surface. However, the mobility of LHCII protein becomes up to $\sim$ 10 fold slower at acidic pH as shown by [3]. Also, as the size of the clusters gets bigger, the lateral diffusion of the aggregates becomes even slower restricting their mobility. This is explained by Saffman-Delbruck equation, $D \propto ln(1/R)$, with $D$ and $R$ as the diffusion constant and the radius of the protein assembly [16]. Once these factors are taken into account, the longer time (> 10 hrs.) required by the system to reach an equilibrium is justified. A more quantitative description of the kinetics of cluster formation in liposomes is given in Sec. 6.

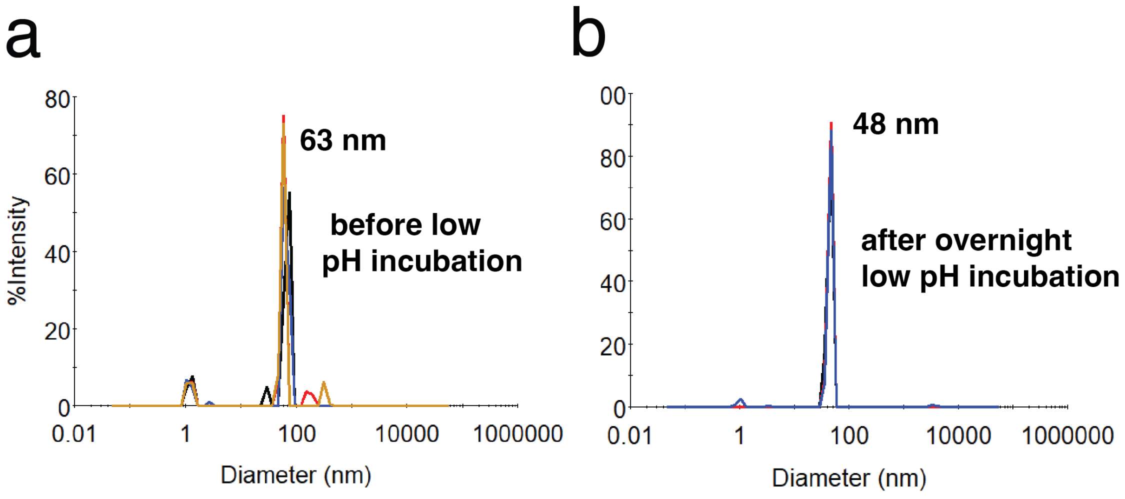

Figure S10: **pH-dependent measurements of proteoliposome.** Size of the proteoliposome (<N>=5) sample (a) before and (b) after the overnight incubation as obtained by dynamic light scattering (DLS) measurements. This indicates that the liposomes remain intact after incubation. The slight difference in diameter may be due to a change in the viscosity of the solvent.

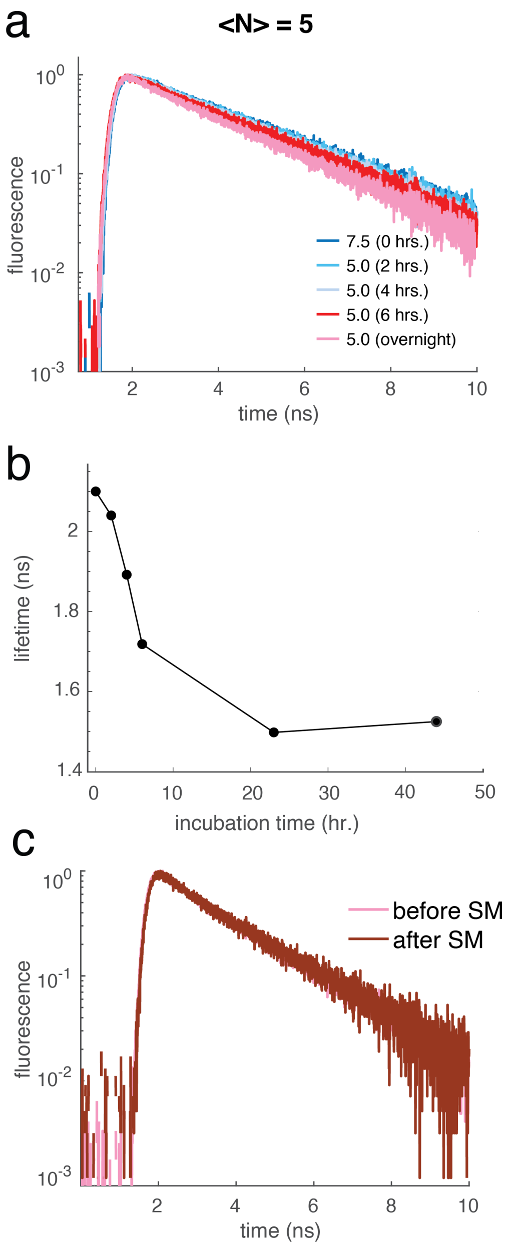

Figure S11: **pH-dependent measurements of proteoliposome.** (a,b) Fluorescence lifetime decays and the average lifetimes of the sample at the different stages of low pH incubation. (c) Single-molecule measurements of the incubated samples were made after ∼10 hrs. of incubation. The overlay of the fluorescence decays of <N>=5 sample before and after the single-molecule measurements shows that the average lifetimes remain intact during the measurements.

### 6 Kinetics of Cluster Formation in Liposomes

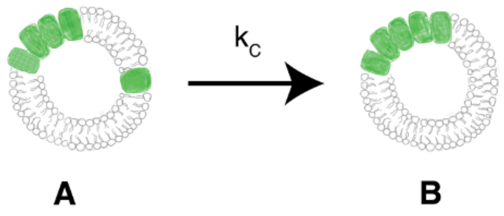

Figure S12: Clustering of proteins from configuration A to B happens at a rate of $k_c(1/\tau_c)$.

Let's assume there are five proteins on the liposome surface with a radius $R$. What is the rate of conversion from configuration A to configuration B as shown in Figure S12? The lateral diffusion coefficient of LHCII on the lipid surface is $D$. Considering a Brownian motion of LHCII on a liposome which is true in a protein-poor system like this, at a certain time period of Δt, the mean area the proteins travel is $4D\Delta t$. Therefore, the fraction of the surface covered by the protein ($f$) at this time interval is,

$$f = \frac{4D\Delta t}{A} = \frac{4D\Delta t}{4\pi R^2} = \frac{D\Delta t}{\pi R^2} \tag{S1}$$

where $A$ is the surface area of the liposome.

Now, the probability that within this random walk, the single LHCII trimer comes close to the other cluster of four proteins and quenching happens is:

$$\frac{D\Delta t}{\pi R^2} * p \tag{S2}$$

where $p$ is the encounter probability. Here, we assume that the diffusion coefficient of the bigger cluster is slow and therefore it remains relatively immobile compared to the single LHCII.

During this quenching event, the concentration of unquenched configuration ($C$), decreased by an amount of $\Delta C$.

Therefore,

$$\frac{\Delta C}{C} = -\frac{D\Delta t}{\pi R^2} * p \tag{S3}$$

At the limit of $\Delta t \leftarrow 0$, the above equation becomes,

$$\frac{dC}{C} = -\frac{D * p}{\pi R^2} dt \tag{S4}$$

Integration of the above equation gives,

$$C = a * exp(-\frac{t}{\tau_c}) + b \tag{S5}$$

where, $\tau_c = \frac{\pi R^2}{D*p}$ is the time constant of cluster formation. a and b are constants.

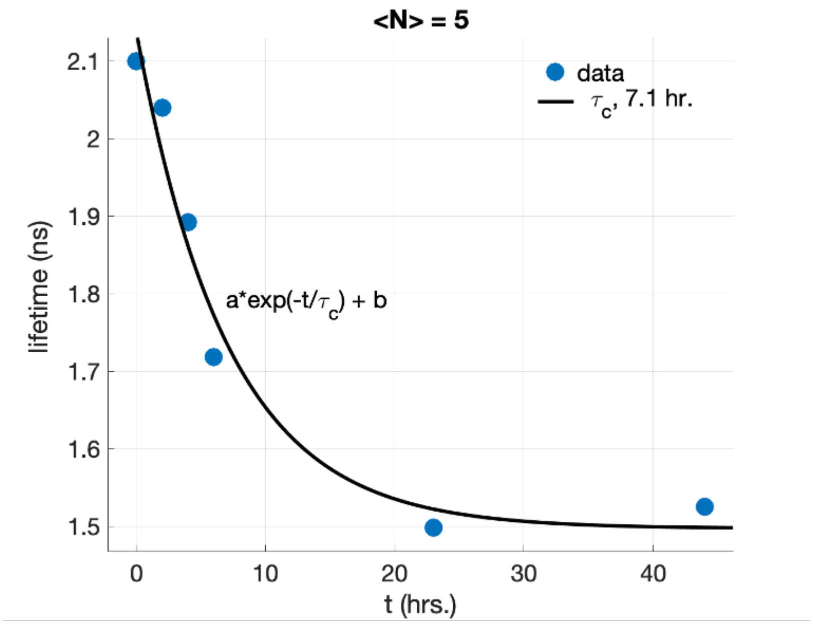

Figure S13: Lifetime drop in the sample with <N> = 5 sample with liposome diameter of 50 nm upon incubation at pH 5. The blue-filled circles are experimentally obtained lifetimes and the black line is the exponential fit.

Figure S13, shows the drop in lifetime for <N> sample as a function of incubation time obtained experimentally. A single-exponential fitting of the trace gives a $\tau_c$ of 7.1 hrs. Again, the encounter probability for two proteins on the liposome size of 25 nm is given as p = 0.006 (ref. analytical solution of cluster formation, Sec 12). The above results give us a diffusion time constant of LHCII under low pH:

$$D = \frac{\pi R^2}{\tau_c * p} = 1.3 \times 10^{-17} m^2/s \tag{S6}$$

The above results assume that the drop in lifetime for <N> = 5 sample is largely from the clustering of configuration A to B (Figure S12). This is approximately true as considering the LHCII-LHCII interaction energy at neutral and low pH as -5.4 and -7.2 $k_B T$, the population of configuration B changes from 23% to 74%. For more rigorous estimation, the contributions from all other configurations and their quenching events need to be accounted for. It is notable that the diffusion constant obtained here $\sim$ is 100-fold slower than that reported from FRAP measurements [7]. This could be due to the slow mobility of LHCII at low pH due to hydrophobic mismatch as shown here by MD simulation [3].

Using the above analysis, it is straightforward to show that with bigger liposomes, the time constant for cluster formation is very slow. For instance, with a liposome of radius 50 nm ($R$=50 nm) and <N> = 5,

$$\tau_c = \frac{\pi R^2}{D * p} = 129 hrs. \tag{S7}$$

Here, the encounter probability, p, is used as 0.0013 (ref. analytical solution, Sec 12).

For proteoliposomes with higher protein density, the analysis of cluster kinetics is complex because it involves multiple configurations. For instance, in proteoliposomes with <N>=10, the 42 different configurations get clustered at different extents upon low pH incubation. However, only accounting for the conversion of the configuration of 9 clustered and one free LHCII to a configuration where all 10 LHCII are clustered, the time constant of cluster formation is estimated as $\sim$ 50 hrs. Therefore, to observe the steady state drop in the lifetime of the <N>=10 samples upon low pH incubation, its needs to be incubated at least $3 \times 50 = 150$ hrs. considering exponential kinetics. However, LHCII-proteoliposomes are not stable for that many hours particularly at low pH as revealed by a reduction in their size observed in dynamic light scattering. Therefore, the low pH measurements of proteoliposomes with protein densities higher than five could not be measured reliably in this work.

### 7 Single-molecule Measurements

The protocol for immobilization of LHCII-proteoliposome samples on biotinylated coverslip has been described in the main text.

For the single-molecule measurement of immobilized samples, the excitation was generated by a tunable fiber laser (FemtoFiber pro, Toptica Photonics, 80 MHz repetition rate, 130 fs pulse duration, 610 nm, 4 nm full-width half maximum (fwhm)), passed through a pinhole, and directed into a home-built confocal microscope. The excitation was focused by an oil-immersion objective (UPLSAPO100XO, Olympus, NA 1.4) onto the samples immobilized on a coverslip. The coverslip was mounted on a piezostage controlled by a home-written Labview-based software. The stage was used to raster-scan on a 5µm x 5µm area to detect single particles (Figure S14).

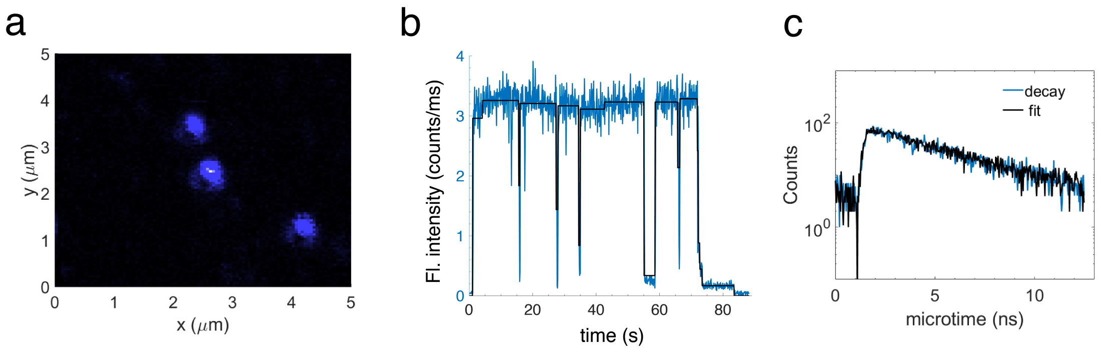

Figure S14: **Single-molecule measurement of LHCII-proteoliposome.** (a) A representative confocal image showing three LHCII proteoliposomes immobilized on a coverslip. (b) A typical fluorescence trace upon excitation of a single LHCII proteoliposome. The regions of constant fluorescence intensity or 'states' are identified by a change-point finding algorithm. (c) A Representative fit for the lifetime decay of a 'state' in single-molecule experiments based on maximum likelihood estimation (MLE) algorithm.

.

The concentration of the sample was diluted to $\sim$4-6 LHCII-proteoliposomes per 25 µm$^2$ scanned area. Subsequently, the laser was parked on the detected spots. The emission of the sample was collected through the same objective and separated from the excitation using a dichroic (ZT647rdc, Chroma) and bandpass filters (ET700/75m, Chroma and and ET690/120x, Chroma). The laser spot size was 280 nm - 380 nm (fwhm) (Figure S8 b). The average power and intensity of excitation used for the single-molecule measurements were 350 nW and 5250 nJ/cm$^2$ per pulse, respectively. Emission was detected by a silicon-based single photon counting avalanche photodiode (SPCM-AQRH, Excelitas Technologies). A time-correlated single-photon counting (TCSPC) module (Time Tagger 20, Swabian Instruments) was used to record the macrotime and microtime for each detected photon [8]. The instrument response function (IRF) measured from the scattered signal was $\sim$ 400 ps (fwhm). Fluorescent traces from >100 complexes were recorded for each set of measurements.

Table S3: Single-molecule lifetime of LHCII in liposome at various <N>. The values in the parenthesis are the errors obtained from standard deviations of the bootstrapped distributions.

| protein:lipid | <N> | $\tau_{median}$ (ns), pH=7.5 | $\tau_{median}$ (ns), pH=5.0 |
|---|---|---|---|
| 1:7000 | <1 | 2.29 | 2.02 (*0.02*) |
| 1:5000 | 1 | 2.45 (*0.04*) | - |
| 1:2500 | 2 | 2.27 (*0.08*) | 1.75 (*0.02*) |
| 1:1000 | 5 | 1.47 (*0.04*) | 0.96 (*0.02*) |
| 1:880 | 6 | 1.53 (*0.04*) | - |
| 1:510 | 10 | 1.22 (*0.02*) | - |

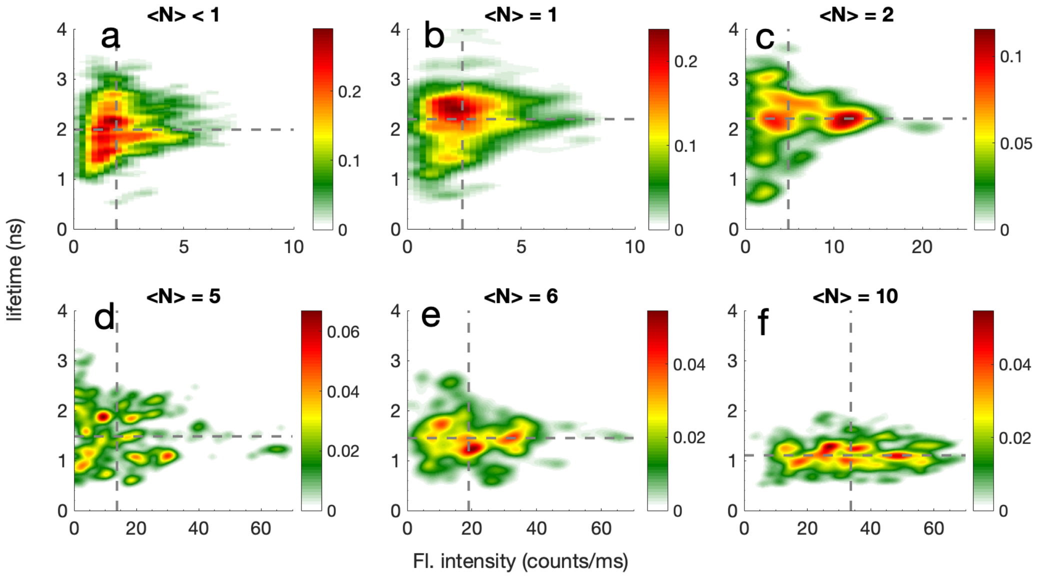

Figure S15: 2D density plots displaying lifetime and fluorescence intensity of LHCII-proteoliposome samples with an average number of proteins (<N>) ranging from less than one to ten. The dotted lines show the median lifetime and fluorescence intensity of the samples. The vertical color bars display probability densities.

### 8 Single-molecule Data Analysis

**Lifetime fits by maximum likelihood estimation (MLE).** The TCSPC module recorded the absolute arrival time of the photons to the detector (macrotime) and the time relative to a trigger pulse (microtime). The macrotime was binned at 100 ms resolution to generate fluorescence intensity traces. Regions of constant intensity (states) were identified by a change-point algorithm [22]. The threshold for the change-point algorithm was adjusted manually based on the inspection of fluorescence traces of the individual complexes. The microtimes of the photons from each state were binned at 280 ps resolution and the resultant histogram was fit with a single-exponential function convolved with the IRF to obtain excited state lifetimes. States with fewer than 500 photons were removed from the analysis as reliable fitting of a lifetime could not be obtained. Due to the low signal-to-noise (S/N) ratio encountered in single-molecule measurements, the decay curves were fit by an algorithm based on maximum likelihood estimation (MLE). MLE captures the inherent Poissonian noise of the single-molecule photon detection and provides robust fitting of the data compared to the conventional non-linear least square method in the low S/N regime [10]. The details of the fitting are provided in [9].

### 9 Density- and pH-induced lifetime drop in LHCII proteoliposome.

In recent years, the aggregation of LHCII has been explored in model membranes including liposomes and nanodiscs [1, 11, 18, 19]. It was shown that with the increase of protein density in LHCII-proteoliposome, the excited state lifetime of the LHCII complexes shortened from $> 3$ ns (in the detergent) to less than 1 ns. This establishes that LHCII proteins have a very strong tendency to aggregate even in the absence of a pH gradient. Here, we also observed similar protein density-dependent quenching of LHCII in the liposome as described in the literature [1, 11, 19]. We define the density-induced lifetime drop ($\Delta\tau_{density}$) in the following way:

$$\Delta\tau_{density} = \frac{\tau_N - \tau_1}{\tau_1} \times 100 \tag{S8}$$

where, $\tau_N$ and $\tau_1$ are the lifetimes of proteoliposome with average protein densities of $N$ and 1, respectively. We observe a $\sim 30\%$ density-induced drop in ensemble lifetime once the average proteins in liposome are increased from less than one to 10 (Figure 1d main text, Table S2). A similar trend is observed in the lifetimes obtained from single-molecule measurements (Table S3).

It has been shown in the intact chloroplast [5] and in the model membranes [13, 18] that lowering the pH enhances fluorescence quenching. Nicol et al. [13] reported that in the LHCII-packed liposomes (lipid:protein = 100:1 or 131:1), lowering the pH from 7.5 to 5.5 shortens the lifetime by 20-30%. Here, we define the pH-induced lifetime drop ($\Delta\tau_{pH}$)as the following way:

$$\Delta\tau_{pH} = \frac{\tau_{N,7.5} - \tau_{N,5.0}}{\tau_{N,7.5}} \times 100 \tag{S9}$$

where, $\tau_{N,7.5}$ and $\tau_{N,5.0}$ are the lifetimes of the LHCII proteoliposome with average protein densities of $N$ at pH 7.5 and 5.0, respectively.

### 10 Clustering of interacting proteins.

When a certain number of proteins with pairwise attractive potential are allowed to interact with each other by restricting them to a finite surface area they tend to form clusters. The driving force of this cluster formation is the reduction of free energy. Based on the mode of clustering, i.e., how the proteins in the cluster are connected, we can classify different types of configurations from these interacting proteins. The number of possible type of configurations for $N$ number of proteins can be obtained by solving the equation below [17]:

$$\sum_{l=1}^{N} l * m_l = N \tag{S10}$$

where, the system forms a cluster with $m_1$ number of unit cluster, $m_2$ number of cluster of 2 proteins, .... $m_l$ number of cluster of l proteins and so on.

For instance, with $N = 5$, there are seven solutions that satisfy the above equation (Eqn. S10). In other words, seven different configurations are possible as shown in Figure S16a. These seven solutions of the equation S10 are displayed in Table S4. The first solution ([0 0 0 0 1]) represents the configuration where all the proteins form a cluster while the last solution ([5 0 0 0 0]) represents where all the proteins remain unclustered. The rest of the five configurations consist of various numbers of clustered and free proteins.

Table S4: Solutions of the Equation S10 with N = 5. The resultant **m**-matrix represents all the possible cluster configurations for this $N$ with their geometric connectivity.

| $m_1$ | $m_2$ | $m_3$ | $m_4$ | $m_5$ |
|---|---|---|---|---|
| 0 | 0 | 0 | 0 | 1 |
| 0 | 1 | 1 | 0 | 0 |
| 1 | 0 | 0 | 1 | 0 |
| 1 | 2 | 0 | 0 | 0 |
| 2 | 0 | 1 | 0 | 0 |
| 3 | 1 | 0 | 0 | 0 |
| 5 | 0 | 0 | 0 | 0 |

As the number of proteins increases, the possible number of configurations also increases. For example, with 10 proteins, 42 configurations are possible. Figure S16b displays the number of configurations as a function of the number of proteins. The number of possible configurations is obtained by solving Eqn. S10.

In this work, we prepared LHCII-proteoliposomes with various lipid-to-protein ratios that generate liposomes with an average number (<N>) of proteins up to 10. For a given number of LHCII, the equilibrium population of a certain configuration ($p$) arising from the clustering depends on both the probability of forming the configuration ($w$) and the enthalpy ($H$) weighted by the Boltzmann factor. This can be expressed as follows:

$$p = w * e^{-H/k_B T} \tag{S11}$$

where, $k_B$ and $T$ are Boltzmann constant and temperature, respectively. In the following section, we discuss the way we compute the equilibrium population ($p$) of these configurations. The populations are then exploited to extract the pairwise interaction energies ($J$) of LHCII proteins.

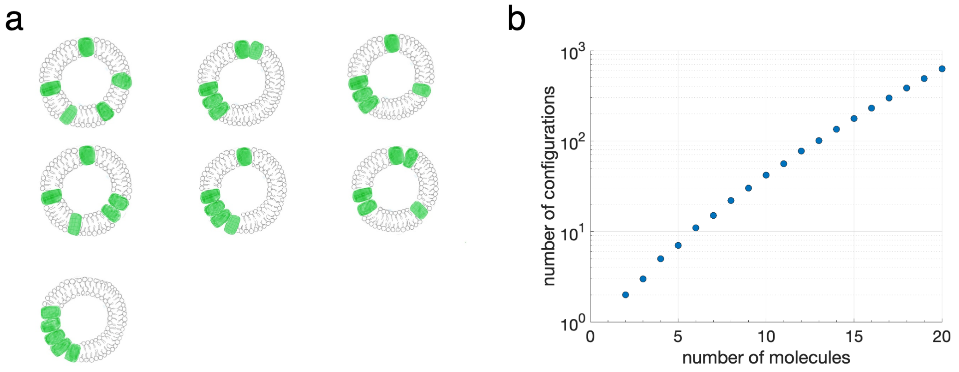

Figure S16: (a) Cartoons of the seven different configurations of clusters possible when five LHCII proteins are incorporated into a liposome. (b) The number of possible configurations increases with the number of proteins.

### 11 Determination of the equilibrium population of cluster configurations from simulation.

To determine the probability of forming different configurations ($w$), we use Monte-Carlo simulations to randomly distribute the proteins on the liposome surface. A homemade Matlab (Mathworks, Inc.) code was used for this. The liposome is modeled as a spherical surface with a radius, $R$, of 25 nm. Proteins (LHCII) are randomly assigned $x$, $y$, and $z$ coordinates between 0 and 1. These coordinates, $C$, are then projected onto the surface of this sphere using the equation:

$$C' = \frac{R.C}{\sqrt{C.C}} \tag{S12}$$

The distance, d, between proteins is then calculated by:

$$\theta = cos^{-1}(\frac{C'_1.C'_2}{R^2}) \tag{S13}$$

$$d = R\theta \tag{S14}$$

where $\theta$ is the angle between the two proteins.

If any two proteins are separated by a distance of less than 6.5 nm (size of LHCII trimers), the configuration is automatically thrown out and a new random configuration is chosen. This is to avoid any non-physical overlap of two proteins on the liposome surface. Any proteins separated by a distance of 6.5-7.5 nm are assumed to be in the same cluster and able to interact. The number of clusters and the number of proteins in each cluster determines the configuration. Each configuration has an assigned enthalpy calculated by:

$$H = \sum_{i=1}^{N} J * (n_i - 1) * m_i \tag{S15}$$

Where $J$ is the pairwise interaction energy, $n_i$ is the number of proteins in the cluster of size $i$, $m_i$ is the number of clusters of size $i$, and $N$ is total the number of proteins in the liposome.

Table S5 displays the probability of different configurations with the number of LHCII in liposomes from 2 to 6. The simulations are typically run for $10^7 - 10^9$ number of iterations until the population of various configurations converges. All of the possible configurations are recovered with these iterations for N up to 5. However, for the systems with molecules beyond 6, all configurations are not observed even after running the iterations up to $10^9$ times. As it is displayed in Table S5, for $N = 6$, the configuration where all the LHCII molecules clustered (configuration $a$) is very unlikely just considering the $w$ factor. However, to properly account for the equilibrium population of the configurations, probabilities of all the configurations are necessary. This is because some configurations might have a low probability of occurrence just considering the entropy factor but could be possible when the enthalpy factor is also accounted for. Therefore, we solved the probability of the cluster configurations via an analytical approach as described below.

Table S5: The probability ($w$) of different configurations as computed from randomly distributing LHCII into liposome surface. The number of iterations used in the simulations is $10^7 - 10^9$.

| configuration | N =2 | N=3 | N=4 | N=5 | N=6 |
|---|---|---|---|---|---|
| a | 0.006 | $6 \times 10^{-5}$ | $1.2 \times 10^{-6}$ | $2.0 \times 10^{-8}$ | $<1.3 \times 10^{-9}$ |
| b | 0.994 | 0.017 | $99 \times 10^{-6}$ | $3.6 \times 10^{-6}$ | $4.9 \times 10^-8$ |
| c | | 0.98 | $25 \times 10^{-5}$ | $5.5 \times 10^{-6}$ | $1.0 \times 10^{-7}$ |
| d | | | 0.033 | 0.0005 | $2.7 \times 10^{-6}$ |
| e | | | 0.97 | 0.0006 | $1.5 \times 10^{-7}$ |
| f | | | | 0.055 | $2.2 \times 10^{-5}$ |
| g | | | | 0.94 | 0.0014 |
| h | | | | | 0.0012 |
| i | | | | | 0.080 |
| j | | | | | 0.92 |

#### 12 Analytical solution of the probability of cluster formation on liposome.

Let us consider a liposome with a radius of $R$. The total surface area of the liposome is $4\pi R^2$. Let us also assume that there is a protein molecule with radius $r$ embedded on this liposome surface. Now, what is the probability that if we insert another protein, those two will form a cluster? For this, we need to define the cluster. In this work, we define that if the second molecule is within a distance of $d$ away, we consider those two proteins to form a cluster. Now, as the second protein cannot occupy the space of the first protein, the probability of forming this cluster is proportional to $\pi(r + d)^2 - \pi r^2$ (the yellow shaded region in Figure S17). Therefore, the probability of forming a cluster of two proteins ($p_2$) is given by,

$$p_2 = \frac{\pi(r + d)^2 - \pi r^2)}{4\pi R^2 - \pi r^2} \times 2 \tag{S16}$$

The denominator in the above equation reflects the total surface area available for the second molecule to be on the liposome surface. The factor of 2 accounts for the exchangeability of these two identical proteins. With $R = 25$ nm, $r = 3.25$ nm, and $d = 1$ nm; the above equation predicts

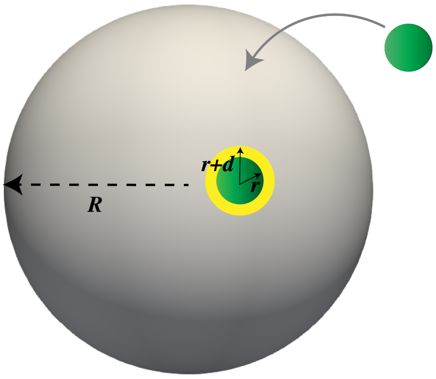

Figure S17: Probability of forming a cluster of two proteins on a spherical surface with radius R is proportional to the area shaded in yellow.

the probability of forming a cluster of two LHCII on the liposome surface is 0.006. This exactly matches the result obtained from the Monte Carlo simulation (Table S5).

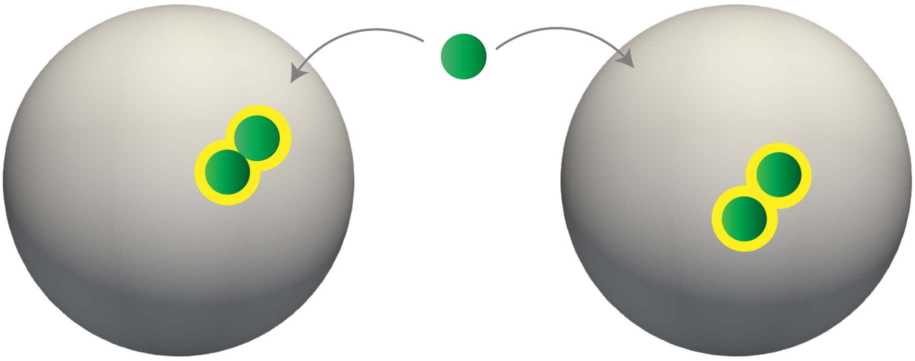

Figure S18: Probability of higher order cluster formation is proportional to the yellow shaded region. Two extreme configurations of clusters are shown in the left (minimum probability/compact arrangement) and right (maximum probability/non-compact arrangement) spheres.

The probability of higher-order cluster formation can be calculated in a similar way. However, for higher-order clusters, different modes of arrangement need to be considered. For example, the probability of cluster formation of three proteins ($p_3$) is proportional to:

$$p_3 = p_2 * p_3' \tag{S17}$$

Where, $p_2$ is the probability of cluster formation of two proteins and $p_3'$ is the probability that the third proteins find this cluster of two proteins on the liposome surface. As similar to discussed above, $p_3'$ is proportional to the yellow shaded regions in Figure 2. Now, when the third molecule is added to the liposome, the cluster of two molecules can be in two extreme arrangements: 1) they can touch each other (Figure S18, left) or 2) they can be in $d$ distance away (as per the definition of a cluster, Figure S18, right). Let us call them compact and non-compact arrangements, respectively. In reality, most of the time, these two proteins in a cluster can be found in between compact and non-compact arrangements.

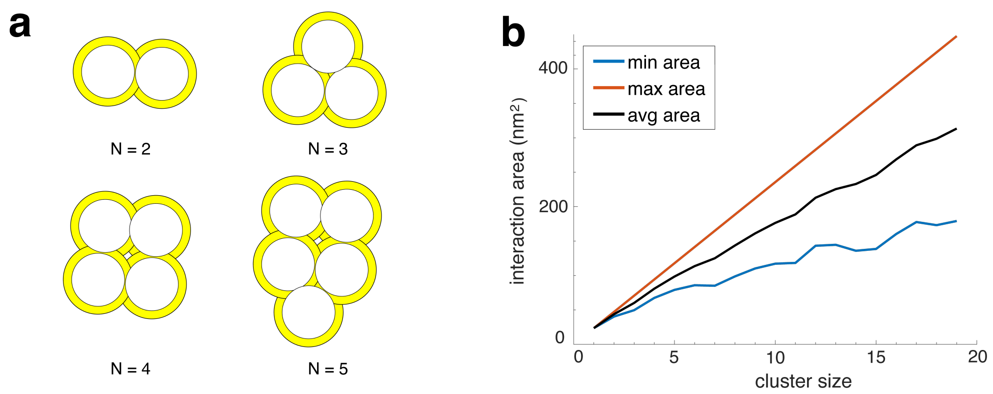

Figure S19: (a) Cluster of different sizes ($N$ = 2,3,4,5) arranged in compact ways. The minimum area available for interactions is shown as a yellow-shaded region. (b) Minimum, maximum, and average area of interactions for different cluster sizes.

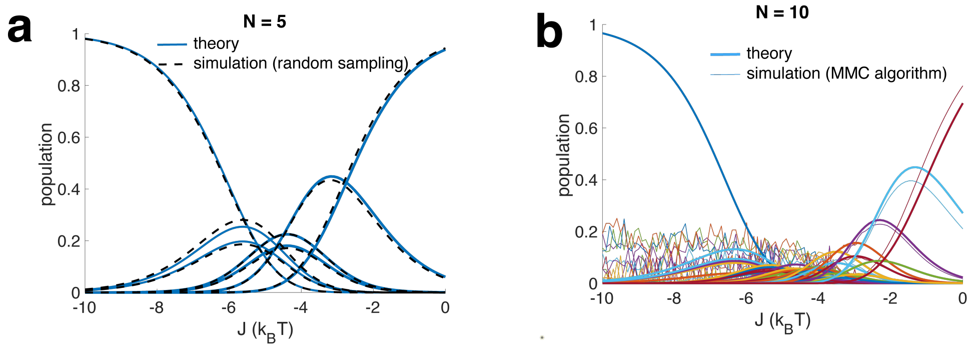

Figure S20: Comparison of the population of different configurations between theory (analytical) vs. simulation obtained using (a) random sampling and (b) MMC algorithms.

Figure S19 a shows the area available for the compact arrangement of clusters having $N$=2-5 proteins. The probability that when another protein added to the liposome would form a cluster is proportional to the yellow-shaded region. This available interaction area for a compact arrangement is the minimum available area. This minimum available area for a number of clusters is computed using a commercial software (`https://www.sketchandcalc.com/`). The maximum available area for the non-compact arrangements for a cluster of $N$ particles is $N * \pi((r+d)^2 - r^2)$. Figure S19 b displays the maximum (non-compact), minimum (compact), and average areas (mean of maximum and minimum area) available for interactions when another molecule/protein is added to the cluster.

This average area ($A_{avg}$) is used to compute the probability of cluster formation. For instance,

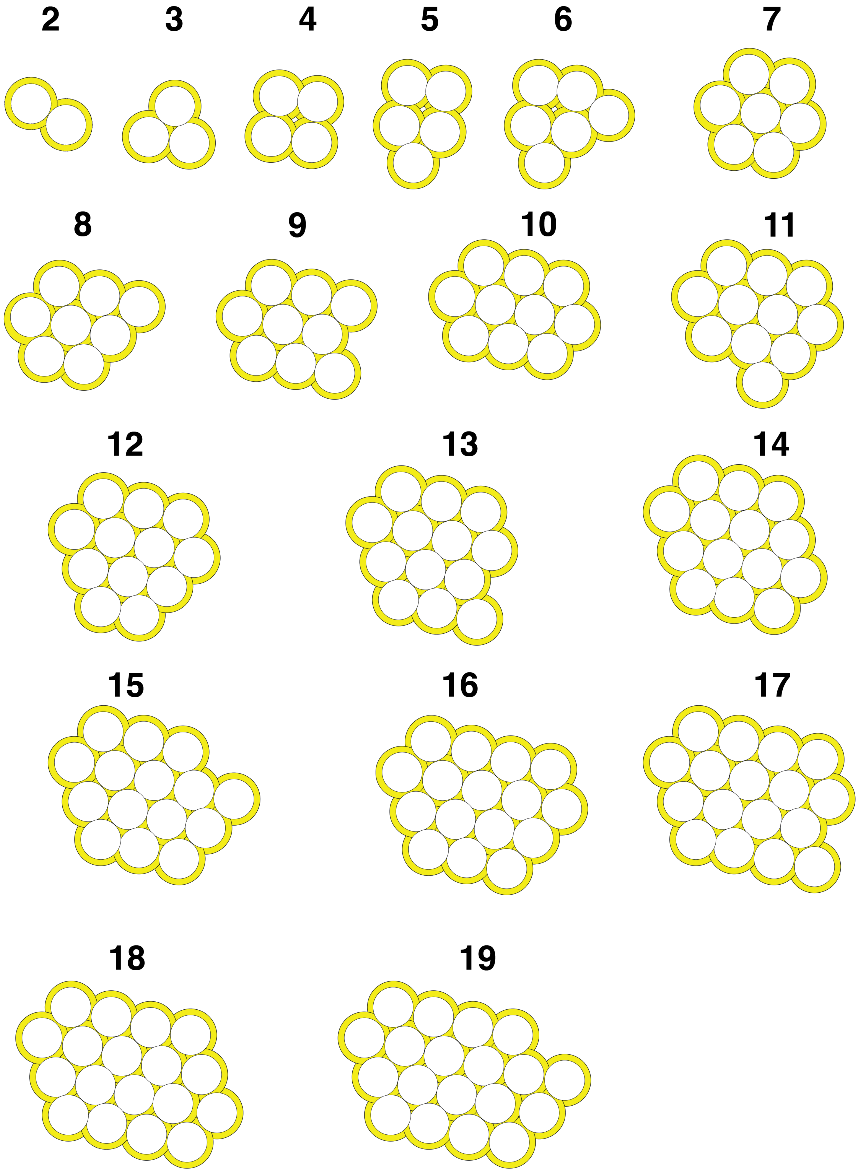

Figure S21: LHCII cluster geometry considered to obtain their analytical probability of formation.

the probability of cluster formation of three proteins ($p_3$) is given as:

$$p_3 = \frac{p_2 * A_{(avg,2)} * 2}{4\pi R^2 - 2 * \pi r^2} \tag{S18}$$

This can be generalized to compute the probability of cluster formation of $N$ protein ($p_N$) as:

$$p_N = \frac{p_{(N-1)} * A_{(avg,N-1)} * 2}{4\pi R^2 - (N-1) * \pi r^2} \tag{S19}$$

Where, $p_{(N-1)}$ and $A_{(avg,N-1)}$ are the probability of cluster formation and the average interaction area available in a cluster of $(N-1)$ number of proteins.

Table S6: Comparison of the probability of the configurations ($w$) obtained from analytical theory and random sampling code.

| N =2 | | N=3 | | N=4 | | N=5 | | N=6 | |
|---|---|---|---|---|---|---|---|---|---|
| theory | sim | theory | sim | theory | sim | theory | sim | theory | sim |
| 0.60 | 0.60 | 0.007 | 0.006 | 1.1e-4 | 1.2e-4 | 2.2e-6 | 2.0e-6 | 5.6e-8 | <1.2e-7 |
| 99.4 | 99.4 | 1.807 | 1.686 | 11e-3 | 9.9e-3 | 4.1e-4 | 3.6e-4 | 4.6e-6 | 4.9e-6 |
| | | 98.18 | 98.31 | 0.027 | 0.025 | 5.3e-4 | 5.4e-4 | 0.95e-5 | 1.0e-5 |
| | | | | 3.615 | 3.346 | 0.05 | 0.05 | 3.3e-4 | 2.7e-4 |
| | | | | 96.34 | 96.62 | 0.07 | 0.06 | 1.3e-4 | 1.5e-4 |
| | | | | | | 6.02 | 5.50 | 2.4e-3 | 2.2e-3 |
| | | | | | | 93.9 | 94.4 | 1.6e-3 | 1.6e-3 |
| | | | | | | | | 0.16 | 0.14 |
| | | | | | | | | 0.13 | 0.12 |
| | | | | | | | | 9.04 | 8.10 |
| | | | | | | | | 90.7 | 91.2 |

**Probability of the configurations with both clustered and unclustered proteins.** Above, we derived the expression of probability to form configurations where all the molecules are clustered. However, if a certain number of proteins are incorporated into liposomes, all the proteins may not form clusters. Some proteins may remain unclustered or free. For instance, if five molecules are incorporated into the liposome, they form a configuration where three of them form a cluster and the other two remain free. What is the probability of forming such configurations? Let us assume a cluster with $m_1$ number of unit clusters, $m_2$ number of clusters of 2 molecules, $...m_l$ number of $l$ molecules, and so on. The probability of forming such a configuration is given by:

$$(\prod_{l=2}^{N} p_l^{m_l}) * \frac{N!}{\prod_{l=1}^{N} m_l!(l!)^{m_l}} \tag{S20}$$

The first part of this equation comes from the joint probability of forming clusters of different sizes, whereas, the second part accounts for the exchangeability of the proteins. Consider the example of the above-mentioned configuration of five proteins with an aggregate of three proteins and two unclustered proteins (unit clusters). The probability of forming such a configuration is given by:

$$(p_3)^1 * \frac{5!}{[2!(1!)^2] * [1!(3!)^1]} = p_3 * 10 \tag{S21}$$

Figure S20 and Table S6 compares the probability of the configurations obtained from the analytical approach (Eqn. S20) and simulations using both random sampling and MMC algorithms. Clearly, there is a good agreement between these two methods. It looks like that although the population obtained from the MMC algorithm agrees reasonably well with theory at $J$ values in the range of -4 $k_BT$ to 0 $k_BT$, it deviates from theory at $J$ = -10 to -4 $k_BT$. The reason behind this may be even after multiple rounds of iterations (∼100 million), heavily clustered states (all clustered) do not appear in the MMC simulations and therefore the contribution of these populations comes out as zero (Eqn. S11). However, as these configurations are energetically favorable, they show up in the population obtained from the analytical theory.

### 13 Photokinetic modeling of cluster-mediated lifetime quenching of LHCII in lipid environment.

Once the chlorophyll molecules of LHCII get excited from the ground state ($S_0$) to the singlet excited state ($S_1$), there can be several pathways that lead to its relaxation to the ground state. A fraction of the population relaxes back to the ground state via radiative transition. Another fraction gets trapped into the Chl triplet state which is readily transferred to the Car triplet state via triplet transfer. At higher illumination intensities when the number of excitations per pulse is greater than unity, singlet-singlet annihilation (S-S annihilation) can occur. On the other hand, at a higher repetition rate of the laser, there can be a significant build-up of the triplet state populations due to their longer microsecond lifetimes. This triplet state can quench the singlet excited state population via singlet-triplet annihilation (S-T annihilation) providing another non-radiative pathway for excited state decay of the S1 state. This essentially shortens the excited state lifetime of chlorophyll molecules further (Figure S22).

The number of excitations per pulse ($n$) of LHCII can be computed as follows:

$$n = \frac{I * \sigma * \lambda}{hc} * \frac{1}{w} \tag{S22}$$

$$I = \frac{P}{\pi * a * b} \tag{S23}$$

$$\sigma = 2.303 * \frac{\epsilon_{610}}{NA} \tag{S24}$$

where, $I$, $\sigma$, and $w$ are laser intensity, absorption cross-section of a LHCII trimer, and repetition rate of the laser (80 MHz) respectively. $P$ is laser power (350 nW). $a$ (280 nm) and $b$ (380 nm) are the laser spot sizes (FWHM). $\lambda$ is the excitation wavelength (610 nm). $\epsilon_{610}$ ($4.36 \times 10^5$ $M^{-1}cm^{-1}$) and NA are the extinction coefficient of a LHCII trimer at 610 nm and Avogadro's constant, respectively. $h$ and $c$ are Planck's constant and speed of light, respectively. Based on the above calculations, the excitation per pulse for a single LHCII is 0.0269, which is well below 1 (Figure S23). Therefore, we can ignore the effect of S-S annihilation as a possible quenching pathway of the $S_1$ population.

Now, assuming S-T annihilation as a quenching pathway in our sample, the decay of the excited population ($[S_1]$) can be described by the kinetic equations below:

$$\frac{d[S_1]}{dt} = G(t) - (k + k_{isc})[S_1] - k_{ST}[S_1][T] \tag{S25}$$

$$\frac{d[T]}{dt} = k_{isc}[S_1] - k_T[T] \tag{S26}$$

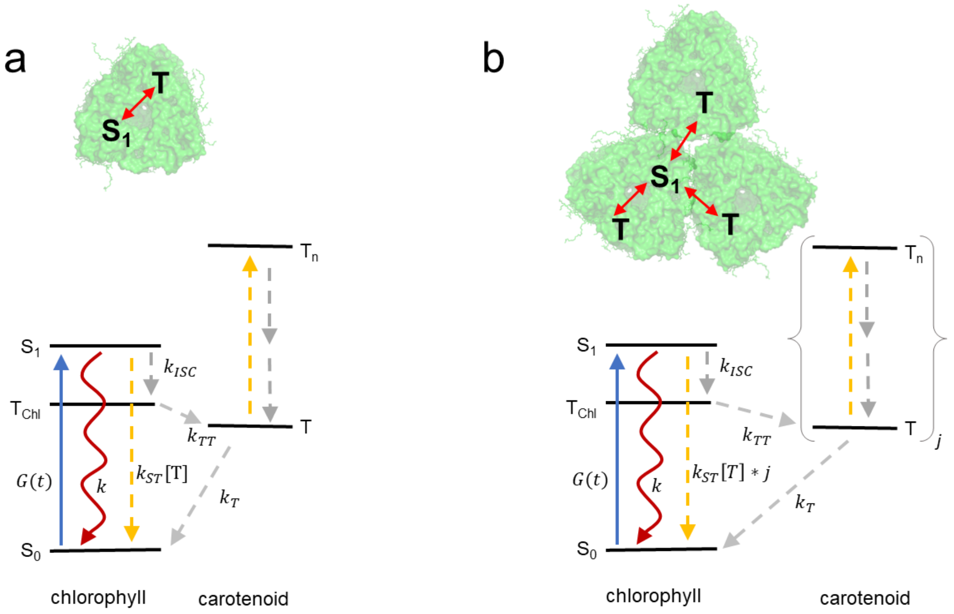

Figure S22: Kinetic model describing excited state relaxation pathways of isolated (a) and clustered LHCII complexes (b). In the clustered complex, the excited state of one LHCII complex can be quenched by the triplet states from other complexes via S-T annihilation process resulting in a faster decay.

Here, $k_{isc}$ (1/8.54 ns$^{-1}$) and $k_T$ (1/7 $\mu s^{-1}$) are rate constants for inter-system crossing and triplet state decay, respectively. $k$ (1/5.81 ns$^{-1}$) is the linear de-excitation rate from $S_1$ state. $k_{ST}$ (1/36 ps $^{-1}$) is the rate constant for S-T annihilation. $G(t)$ is the excitation rate which depends on the absorption cross-section of the sample, illumination intensity, and the repetition rate of the laser. As $T_{Chl}$ readily transfers the population to the triplet state of carotenoids ($T$), this triplet transfer process is effectively eliminated from the above treatment. All the parameters are taken from Gruber et al. [4].

As $k$ and $k_T$ differ by three orders of magnitude, the change in triplet concentration in the steady-state regime between two subsequent laser pulses is almost negligible compared to the accumulated triplet concentration. Therefore, the kinetics of $[S_1]$ is governed by the equation below:

$$\frac{d[S_1]}{dt} = G(t) - (k + k_{isc} + k_{ST} * [T]_0)[S_1] \tag{S27}$$

Where, $[T]_0$ is the population of the triplet state in quasi-stationary condition. Hence, the excited state lifetime of the sample is given by:

$$\tau = \frac{1}{(k + k_{isc} + k_{ST} * [T]_0)} \tag{S28}$$

In this quasi-stationary regime (i.e., at $\frac{d[T]}{dt} = 0$), the population of triplet state can be calculated

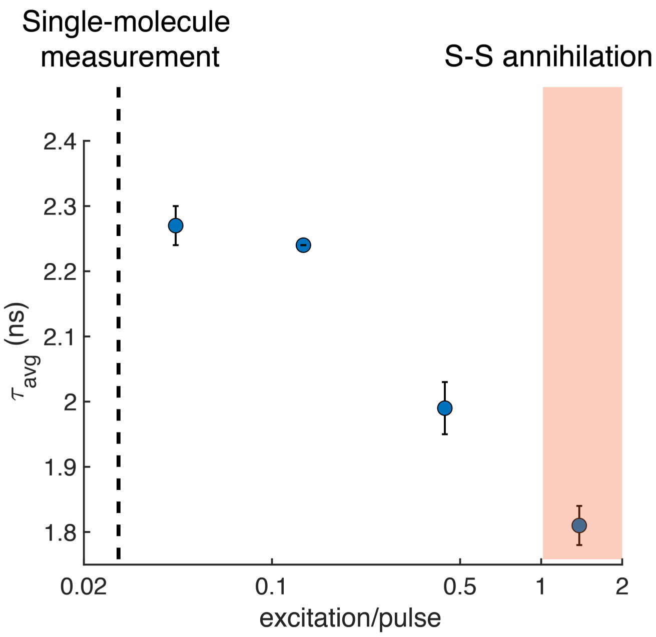

Figure S23: Average lifetime of LHCII proteoliposome with <N> = 5 at pH 7.5 as a function of excitation/pulse. The vertical dotted line shows the excitation/pulse used for single-molecule measurements. The orange-shaded region demonstrates the excitation/pulse > 1 where singlet-singlet annihilation is prominent. Error bars are the standard deviation.

as [23]:

$$\begin{aligned}[T]_0 &= \frac{k_{isc} * [S_1]_0}{k_T} \\ &= \frac{k_{isc}}{k + k_{isc} + k_S T[T]_0} * (\frac{n_0}{k_T}) * w\end{aligned} \tag{S29}$$

Where, $n_0 = \int G(t)dt$ is the fraction of the population into the $S_1$ state after single-pulse. From the above equation, $[T]_0$ is given as:

$$[T]_0 = \frac{-k + \sqrt{k^2 + \frac{4k_{ST} * k_{isc} * n_0 w}{k_T}}}{2k_{ST}} \tag{S30}$$

Combining Eqn. S28 and S30, we get,

$$\frac{1}{\tau} = \frac{k}{2} + k_{isc} + \sqrt{\frac{k^2}{4} + \frac{k_S T * k_{isc} * w}{k_T} * n_0} \tag{S31}$$

Eqn. S31 can be used to estimate the excited state lifetime of single LHCII complexes in the presence of S-T annihilation.

**Lifetime quenching in LHCII clusters** Once multiple LHCII complexes are incorporated in the membrane environment, we observe a shortening of lifetime. This quenching of lifetime depends on protein concentration in the membrane. We model this cluster-dependent quenching in LHCII invoking S-T annihilation.

A small cluster of LHCII proteins can be considered a single supermolecule comprising of various accessible energy levels. As the size of these clusters increases, the singlet excited states can come into contact with triplet states from other LHCII molecules. This enhances the probability of S-T annihilation. Therefore, for a cluster consisting of $j$ number of LHCII proteins, the decay of its singlet excited state can be given as:

$$[S_1] = n_0 * e^{-(k+k_{isc}+j*k_{ST}*[T]_0)t} \tag{S32}$$

In other words, the excited state lifetime of its $S_1$ state is:

$$\tau = \frac{1}{k + k_{isc} + j * k_S T * [T]_0} \tag{S33}$$

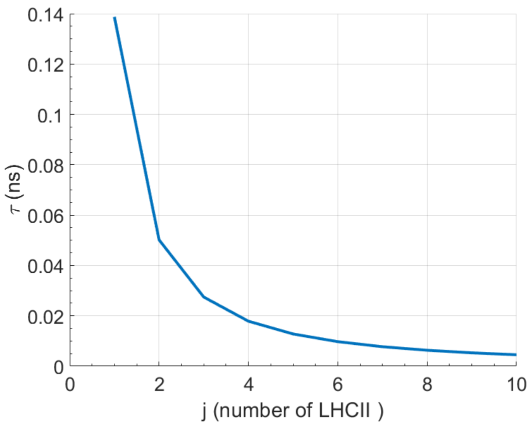

Figure S24: Excited state lifetime of LHCII aggregate as a function of cluster size computed from Eqn. S33.

Figure S24 displays the excited state lifetime of LHCII clusters with cluster sizes from 1-10. As the size of the cluster gets bigger, its lifetime gets shorter due to enhanced S-T annihilation in the bigger clusters.

Now, the computation of excited state lifetime using Eqn. S33 gives a value of 0.14 ns for an isolated LHCII complex. However, this is much shorter than the experimentally observed value of the lifetime of LHCII in detergent ($\sim$ 3.3 ns) or singly-occupied LHCII-proteoliposome (2.8 ns). On the other hand, for a cluster of 10, this model predicts a lifetime of <0.01 ns. This contradicts the experimentally observed lifetime of LHCII-proteoliposome samples with <N> of 10 which is $\sim$ 2.0 ns. This discrepancy is probably due to the fact that in small photosynthetic antenna complexes like LHCII or a small cluster of LHCII contain a limited number of triplet states. Therefore, a simple kinetic model as described above overestimates the contribution of S-T annihilation. To account for the limited number of excitons and their discrete nature, a stochastic model needs to be used as developed in [4].

**Stochastic model of ST annihilation**

Figure S25 demonstrates the stochastic model used to compute the probability of different states ($P_{ij}$). $G(t)$ is the rate of excitation. A rectangular pulse profile was used to model $G(t)$ with

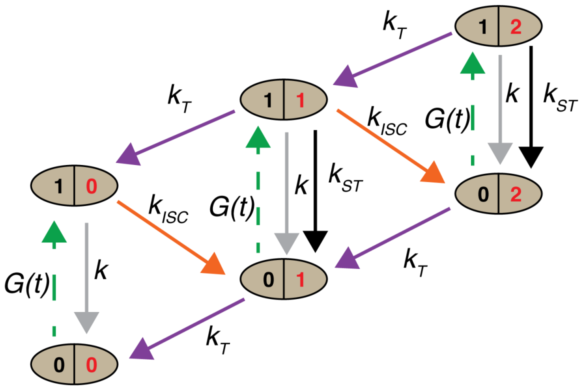

Figure S25: Stochastic model used to compute triplet state population. The ovals represent states with different numbers of singlets (black) and triplets (red) respectively. $G(t)$ is the excitation rate; $k$ and $k_T$ are rate constants for linear de-excitation from singlet and triplet states, respectively. $k_{isc}$ and $k_{ST}$ are the rate constants for inter-system crossing and ST annihilation, respectively. This figure is adapted from [4]

pulse-width and inter-pulse delay of 130 fs and 12.5 ns respectively, similar to the experiments. The probabilities of each state ($P_{ij}$) can be computed by solving the set of coupled differential equations below:

$$\frac{dP_{00}}{dt} = -G(t)P_{00} + kP_{10} + k_T P_{01}$$
$$\frac{dP_{10}}{dt} = G(t)P_{00} - (k + k_{isc})P_{10} + k_T P_{11}$$
$$\frac{dP_{01}}{dt} = k_{isc}P_{10} + (k + k_{ST})P_{11} - (G(t) + k_T)P_{01} + 2k_T P_{02}$$
$$\frac{dP_{11}}{dt} = G(t)P_{01} - (k_T + k + k_{ST} + k_{isc})P_{11} + 2k_T P_{12}$$
$$\frac{dP_{02}}{dt} = -(G(t) + 2k_T)P_{02} + k_{isc}P_{11} + (k + 2k_{ST})P_{12}$$
$$\frac{dP_{12}}{dt} = G(t)P_{02} - (k + 2k_T + 2k_{ST})P_{12}$$

These equations were solved numerically using a home-written MATLAB (Mathworks, Inc.) code with the initial condition, $[P_{00}, P_{01}, P_{10}, P_{11}, P_{02}, P_{12}] = [1, 0, 0, 0, 0, 0]$. The step size of the simulation was 100 fs ($<$ 130 fs, the pulse peak width). The simulation was run up to $w/k_T$ pulses as the triplet states experience that many numbers of excitations by the pulsed laser.

Solutions of the probability states at $t = 1/k_T$ can be used to determine the population of triplet state ($[T]_0$):

$$[T_0] = (P_{01} + P_{11}) + 2 * (P_{02} + P_{12}) = 1.8 \times 10^{-3} \quad \text{(S34)}$$

However, as it is shown in Figure S26a, only $P_{01}$ contribute significantly to the population of the triplet state. The other states have negligible triplet state contributions. This also suggests that a further extended model is not necessary. Figure S26b displays the excited state lifetime of LHCII as a function of their cluster size ($j$). The kinetic parameters used for the simulation are provided in

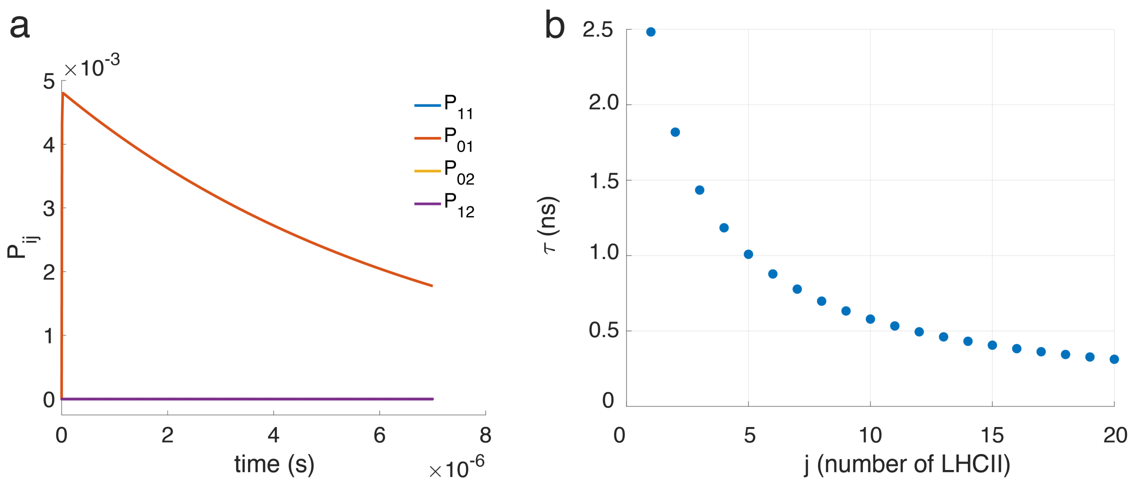

Figure S26: (a) Probability densities of states containing triplet populations as obtained from the numerical simulation. Except for $P_{01}$, the contribution from other states is negligible. (b) Excited state lifetime of LHCII with different cluster sizes ($j$) as obtained from the modeling (Eqn. S33). The kinetic parameters used for the simulation are: $1/k_{ST}$ = 20 ps, $1/k$ = 5.81 ns, $1/k_T$ = 7 $\mu s$, $1/k_{isc}$ = 10 ns.

the caption. Except for the $k_{ST}$, all the other parameters are obtained from Gruber et al. [4]. Here, $k_{ST}$ is adjusted to 20 ps (36 ps reported in [4]) to match the median lifetime observed in isolated LHCII complex ($\sim$ 2.5 ns) under in single-molecule studies [9].

### 14 Intensity dependence of excited state lifetime in the isolated LHCII complexes

The Eqn. S33 shows the excited state lifetime of the LHCII complexes depends on the number of proteins in the cluster ($j$), as well as the stationary triplet state population ($[T]_0$). Now, $[T]_0$ can be increased by increasing the excitation intensity of the sample which in turn should shorten the excited state lifetime. To investigate the validity of this prediction from our model, we measured the single-molecule lifetime distribution of single LHCII complexes immobilized in a PVA matrix at various laser power. The lifetime distributions, the median of these distributions and the prediction from our model are shown in Figure S27. We see a good agreement between the experiment and simulations that validate the model and model parameters.

Table S7: Median excited state lifetime of LHCII (immobilized in a PVA matrix) obtained from the experiment and simulation at different laser power.

| power ($\mu$W) | $\tau_{median}$ (ns, experiment) | $\tau_{median}$ (ns, simulation) |
|---|---|---|
| 0.42 | 2.56 | 2.50 |
| 0.82 | 2.13 | 2.13 |
| 1.6 | 1.96 | 1.78 |

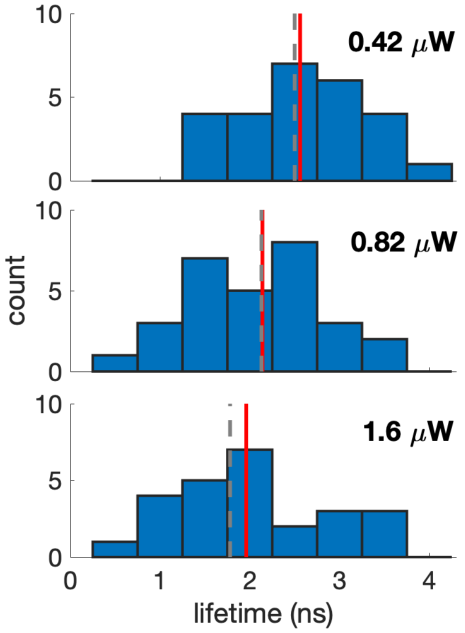

Figure S27: Lifetime distribution of single LHCII complexes immobilized in a PVA matrix at different laser power. The vertical red and dotted grey lines are the median of the distributions from the experiment and the simulation, respectively. The number of complexes recorded for each laser power are: 26, 29 and 25. The lifetime values are demonstrated in Table S7

.

For the measurements in a PVA matrix, LHCII was diluted to $\sim$ 15 pM in a solution containing 5% PVA (MW 78000, 99% hydrolyzed, Polysciences, Inc.), PCA/PCD scavenging mixture and buffer (10mM HEPES, 20 mM NaCl, 0.06% (w/v) GDN, pH 7.5). An argon chamber was used for the single-molecule measurements of LHCII in the PVA matrix to reduce photodegradation [9].

#### 15 Extraction of LHCII-LHCII interaction energy from lifetime data.

Figure S28 demonstrates the steps followed to extract the LHCII-LHCII interaction energy ($J$) from single-molecule lifetime data. Although the clustering reduces the fluorescence intensity of LHCII-proteoliposomes as well as shortens their lifetimes, we employ only lifetime data to extract $J$. This is because the excited state lifetime is a more robust quantity than the fluorescence intensity as it does not get influenced by day-to-day variation of the alignments of our optical set-up and lifetime is independent of illumination intensity. Only the first levels of single-molecule data are used to extract $J$ as in the later levels the proteins undergo photodegradation due to a longer exposure and the quenching model described in the previous section does not include such effects.

Although the average number of LHCII (<N>) embedded in liposomes can be controlled by lipid-to-protein ratio, this number is very heterogeneous [9, 21]. We assume that the number of LHCII

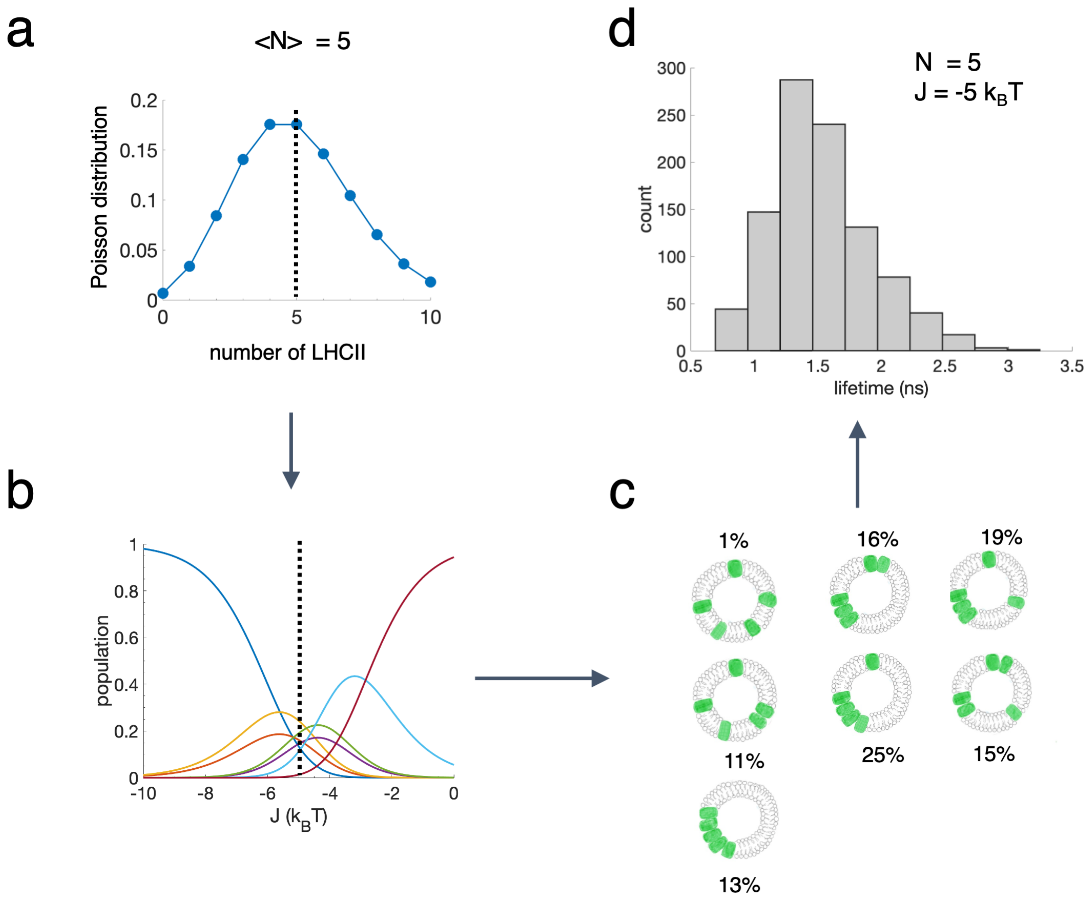

Figure S28: **Extraction of LHCII-LHCII interaction energy from single-molecule lifetime data.** (a) Distribution of the number of LHCII in liposomes for a sample containing on average five LHCII per liposome. In the first step of the simulation, the number of LHCII ($N = 5$ for example) selected based on this Poisson distribution (represented as a vertical dashed line) (b) Population of different configurations for $N$ number of LHCII in liposome at different interaction energies as calculated in Section 11. (c) The relative population of different configurations and the lifetime of clusters based on the quenching model developed in Section 13 are used to compute the average lifetime of the sample at fixed $J$. Steps a-c are run many times to obtain a lifetime distribution for a specific $N$ and $J$.

incorporated in LHCII follows a Poisson distribution. Later, we investigated the effect of Poisson and non-Poisson distribution of LHCII in liposomes on the pairwise interaction energy (Figure S32d). Both analyses show the same LHCII-LHCII interaction energy peaks. Figure S28a shows the Poisson distribution for <N>=5.

In the first step of the simulation, for a sample with a specific <N>, the number of LHCII in a liposome ($N$) is selected from its corresponding Poisson distribution. Once the $N$ is selected, the

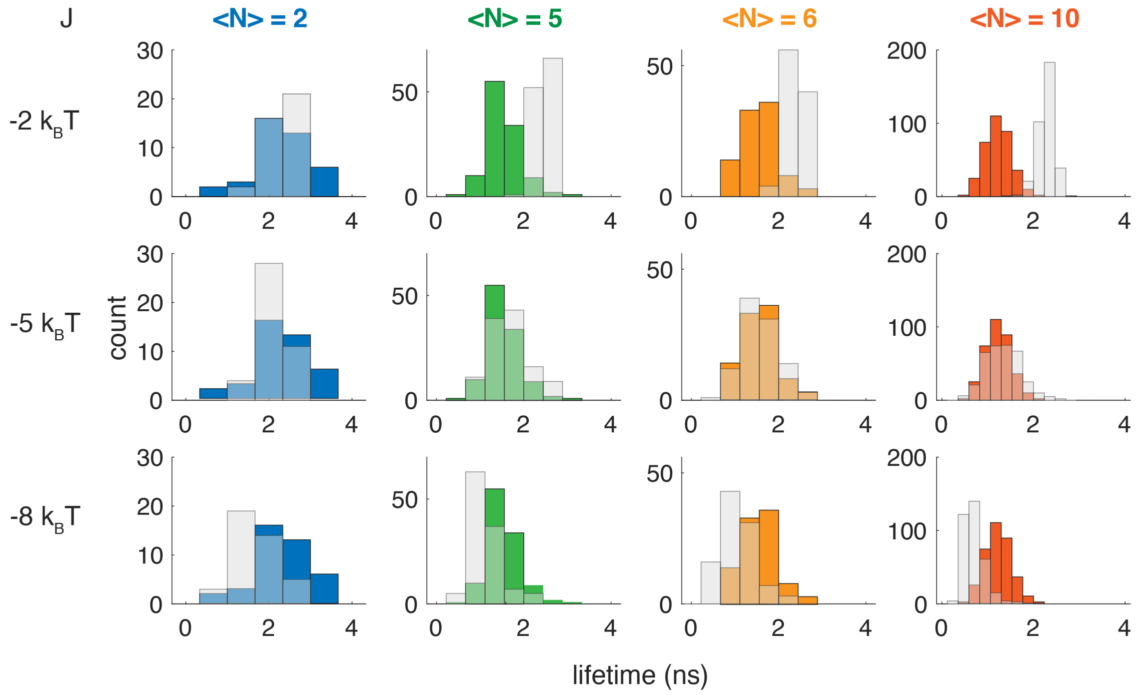

Figure S29: **Comparison between experimental and simulated lifetime distribution at neutral pH**. Lifetime distributions from single-molecule experiments at pH 7.5 are shown in blue, green, orange, and red histograms corresponding to <N> of 2, 5, 6, and 10, respectively. The simulated data (grey histograms) display the lifetime distributions at $J$ = -2, -5, and -8 $k_BT$. The median lifetimes of the distributions are also shown in their corresponding figures.

Matlab program utilizes the population of various configurations at different $J$ (Figure S28b) for that $N$ to compute the average excited state lifetime for these configurations. Based on the size of clusters in a particular configuration, the quenching model predicts their lifetimes (Figure 2a, main text). For instance, in a liposome with five LHCII incorporated the configurations with a cluster of three and a cluster of 2 LHCII, the lifetime ($\tau$) is given as:

$$\tau = \frac{1}{5} * (2 * \tau_{N2} + 3 * \tau_{N3}) \tag{S35}$$

where, $\tau_{N2}$ and $\tau_{N3}$ are the lifetime of clusters consisting of two and three LHCII proteins, respectively. The lifetime of different clusters as a function of sizes is given in Figure 2a (main text) computed from the quenching model (Sec. 13). The steps a-c in Figure S28 are run multiple times to construct a lifetime distribution for a particular LHCII interaction energy (Figure S28d). This figure shows the simulated lifetime distribution for the LHCII-proteoliposome sample with <N> = 5 and $J = -5k_BT$.

Finally, the above procedures are repeated for multiple $J$, ranging from 0 to -10 $k_BT$, and compared with the experimental lifetime distributions. Figure S29 shows the experimental single-molecule lifetime distribution for LHCII-proteoliposomes with <N> of 5, 6, and 10 along with the simulated distributions for $J$ of -2, -5 and -8 $k_BT$. $J$ can be systematically varied to generate lifetime

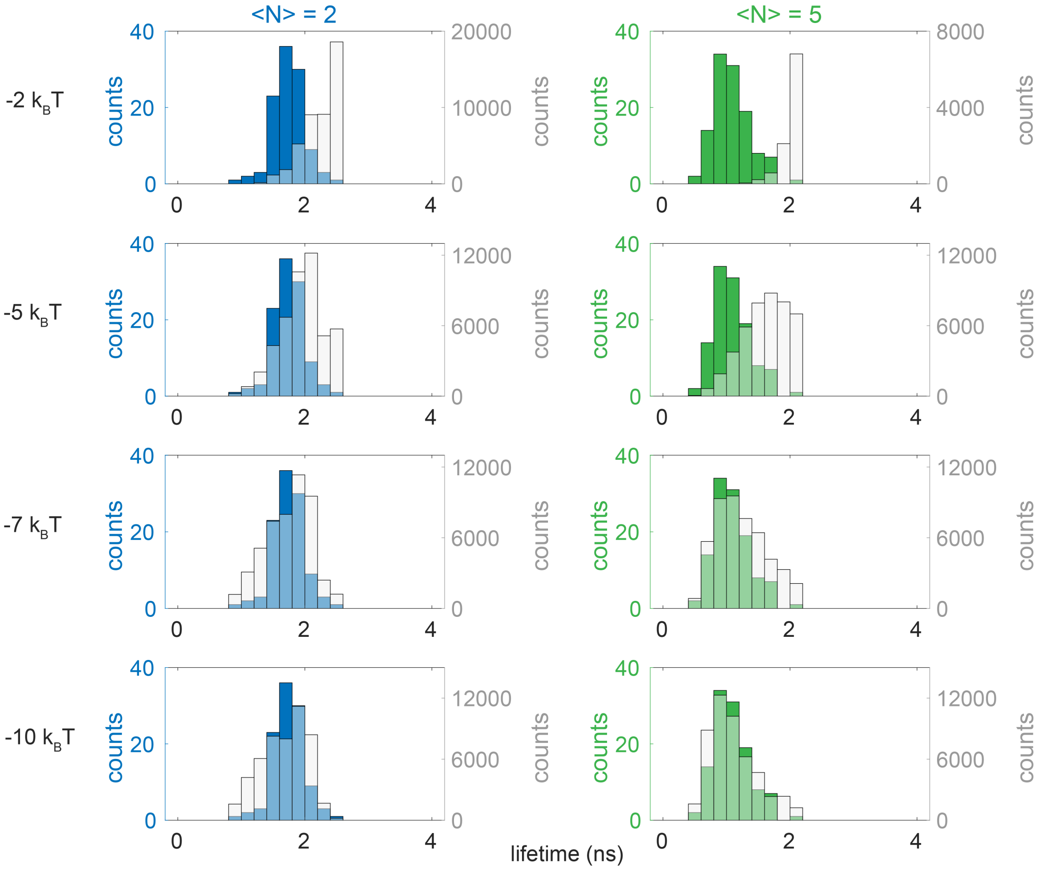

Figure S30: **Comparison between experimental and simulated lifetime distribution at pH 5**. The lifetime distributions from single-molecule measurements are shown in blue and green histograms for <N> = 2 and 5, respectively. The simulated lifetime distributions at $J$ = -2, -5, -7, and -10 $k_BT$ are displayed in grey histograms. At J = ~-7 $k_BT$, the simulated distribution matches well with the experimental ones.

distributions and fit to the experimental data to extract LHCII-LHCII interaction energies. The fits involve converting the lifetime distribution into a probability density function by kernel density estimate (KDE) and subsequent maximum likelihood estimation (MLE) as described below.

Table S8: LHCII-LHCII interaction energies ($J$) obtained from MLE fits. ND = not determined.

| <N> | $J$ pH,7.5 | $J$ pH, 5.0 |
|---|---|---|
| 2 | -5 | -6.6 |
| 5 | -5.6 | -7.4 |
| 6 | -5.2 | ND |
| 10 | -5.4 | ND |
| global | -5.4 | -7.2 |

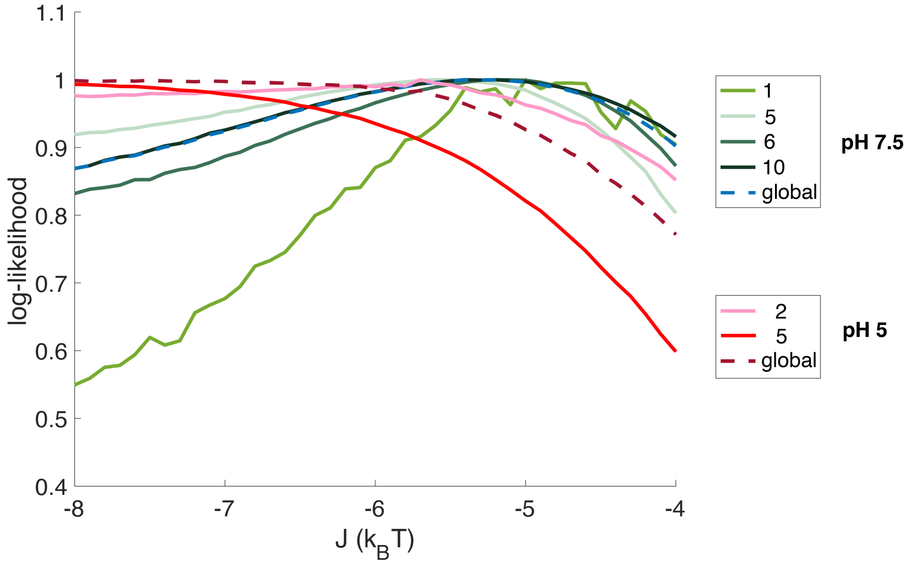

Figure S31: Normalized log-likelihood plots obtained from the MLE fits for the LHCII-proteoliposome samples at different protein densities (solid lines) at pH 7.5 and 5. The corresponding global fits are shown as dashed lines. The peak values of the log-likelihood are demonstrated in Table S8.

**Cross-validated Kernel Density Estimate**. Kernel density estimation (KDE) is a non-parametric method to estimate an underlying probability density function from a data set. We use KDE to estimate the probability density of fluorescent lifetimes as a function of energy. We first simulate 100,000 points at energies ranging from 0 to -10 kBT in 0.1 increments using the method outlined above and then perform KDE on each data set. For an independent, identically distributed set of variables $(x_1, x_2, ..., x_n)$ the KDE is given by:

$$p(x) = \frac{1}{nh} \sum_{i=1}^{i=n} K\left(\frac{x - x_i}{h}\right) \tag{S36}$$

where $h$ is the bandwidth and $K(\frac{x-x_i}{h})$ is the kernel function.

A Gaussian kernel was used in this work. Bandwidth selection is crucial to achieving an appropriate distribution that is neither under nor over-smoothed. We used a method called k-fold cross-validation with k = 5 to pick the bandwidth. In k-fold cross-validation, the data is split into k subsets. One subset is chosen as the test data set while the k-1 subsets are combined into the training set. The KDE is performed using the training data set and then evaluated against the test data set. The process is repeated k times, each time with a different subset used as the test data set. This process was implemented using python's sklearn toolkit [15]. The code is available upon request.

**Maximum Likelihood Estimation**. Maximum likelihood estimation (MLE) is used to estimate which interaction energy best fits the experimental data. The likelihood function is given by:

$$L(\theta) = f(X_1, X_2, .., X_n|\theta) \tag{S37}$$

In words, the likelihood is the probability of observing $x_1, x_2, ..., x_n$ given some parameter $\theta$. The probability of observing a given set of data is:

$$P(X_1 = x_1, X_2 = x_2, ..., X_n = x_n) = \prod_{i=1}^{i=n} p_i \tag{S38}$$

So we can then write:

$$L(\theta) = \prod_{i=1}^{i=n} p_i \tag{S39}$$

In practice, the log-likelihood function is computationally more tractable and is often used.

$$\ln L(\theta) = \ln \left( \prod_{i=1}^{i=n} p_i \right) \tag{S40}$$

$$\ln L(\theta) = \sum_{i=1}^{i=n} \ln(p_i) \tag{S41}$$

The probability density function as determined by KDE is integrated to determine the probability of each experimentally observed lifetime to calculate the likelihood. For the global fit, the likelihood functions of each <N> are added together and the resultant function is optimized to obtain a global value of $J$. Table S8 displays the LHCII-LHCII interaction energies obtained from the global fits and samples with different <N>.

**Parameter dependence.** Polycarbonate membranes with a pore size of 50 nm were used to prepare the liposome. However, the dynamic light scattering (DLS) results of the LHCII-proteoliposome show some variation in their observed sizes (Table S1 and Figure S2). Again, the cluster in the liposome is defined if two proteins are within a distance of $r$ (the yellow shaded region in Figure S32a). In our model, we have assumed this $r$ to be 1 nm. Both the liposome size ($R$) and the interaction distance effect the relative population of different configurations (Eqn. S20) and therefore the extracted LHCII-LHCII interaction energy ($J$). To investigate the dependence of $J$ on $R$ and $r$ used our model, we systematically varied these parameters. Figure S32b and c show the normalized log-likelihood at different $R$ and $r$. We found that these parameters do not affect the extracted $J$ significantly. With $R$ in the range of 25-37.5 nm and $r$ in the range of 0.5-1.5 nm, the extracted $J$ still lies in the vicinity of -5 $k_BT$. Figure S32d displays the dependence of $J$ on the S-T annihilation rate constants. With $k_{ST}$ between 12 and 20 ps, $J$ lies in the range of -5 to -6 $k_BT$. However, at slower $k_{ST}$ ($\sim$ 36 ps), we do not observe a clear peak in the log-likelihood analysis. This indicates, the value of $J$ is weakly dependent on the lifetime of LHCII clusters, which in turn is related to $k_{ST}$ (Eqn. S33). We also investigated the effect of $J$ on assuming that proteins in liposomes follow a Poisson and no distribution (non-Poisson) at all. In both cases, we found a similar peak value in the log-likelihood plot. However, the plot for non-Poisson distribution was a bit noisy and the peak was less pronounced.

#### 16 Surface Charge Calculations

The net surface charge of LHCII ($Q$) was calculated using PROPKA software [14], which takes account of the pKa of the protonatable residues and then applies perturbation factors from the protein including coulomb interactions, desolvation, and electrostatics. The charged surface is then created using the Adaptive Poisson-Boltzmann Solver, which solves the equations of continuum electrostatics [6].

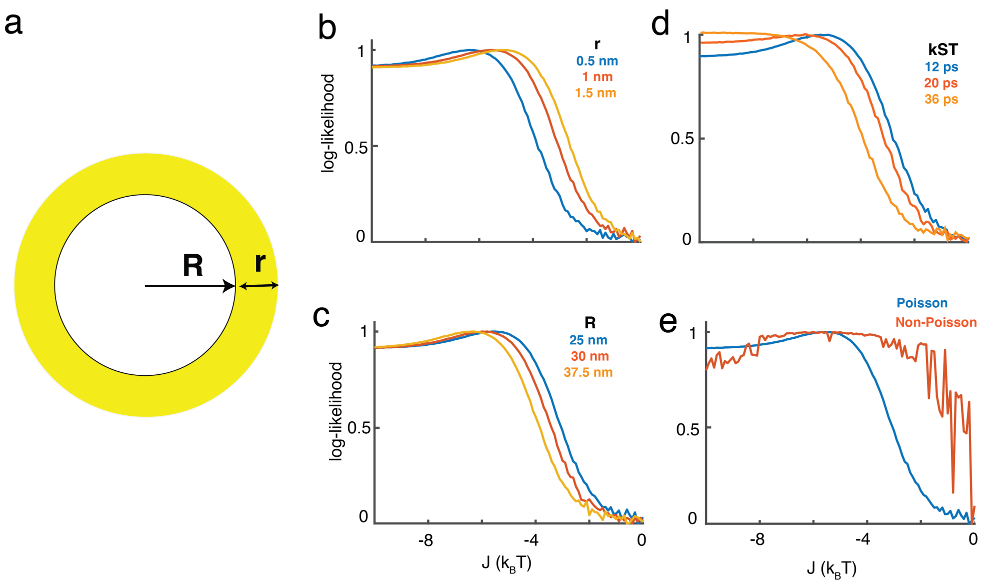

Figure S32: (a) The dependence of LHCII-LHCII interaction energy extracted from our model on the liposome radius ($R$) and interaction distance ($r$) is systematically investigated. Normalized log-likelihood plots at different (b) interaction distances (c) liposome sizes (d) S-T annihilation rate constants and (e) distribution types (Poisson and non-Poisson) are displayed. All the simulations are done for <N>=5.

#### 17 Free energy change in pH-driven clustering in LHCII proteoliposome.

Based on our single-molecule lifetime data and analysis of LHCII clusters in proteoliposome, we observe moderately clustered configurations at neutral pH to heavily clustered ones at low pH. We attribute this to a change in relative configurations at these two conditions. For instance, as Figure S33 shows, the configuration where all the proteins are clustered constitutes 23% of the population. However, at low pH the relative population of this configuration changes to $\sim$ 93%. We aim to quantify the change in enthalpy ($\Delta$H), entropy ($\Delta$S), and free energy ($\Delta$G) of such transition.

Figure S34 shows a schematic representation of how such thermodynamic parameters are calculated. For the calculation of $\Delta$H, the pairwise interaction energies at neutral and low pH are required (Section 15). In this particular case, the total interaction energies are $3J_1$ and $4J_2$. Therefore, $\Delta$H is given as $4J_2 - 3J_1$. On the other hand, $\Delta$S can be estimated from the probabilities ($w$) of those configurations (see Methods, main text).

Now, for the proper calculation of $\Delta G$, the contribution from all other configurations should also be considered. This leads to,

$$\Delta G = \sum_{i=1}^{m} \Delta H_i(J) - k_B T * ln(\sum_{i=1}^{m} \frac{w_{l,i}}{w_{n,i}}) \quad \text{(S42)}$$

where, $\Delta H_i(J)$ is the change in enthalpy associated with $i$-th configurations; $w_{l,i}$ and $w_{n,i}$ are the probabilities of such configurations at pH low and (pH, 5.0) and neutral pH (pH, 7.5), respectively.

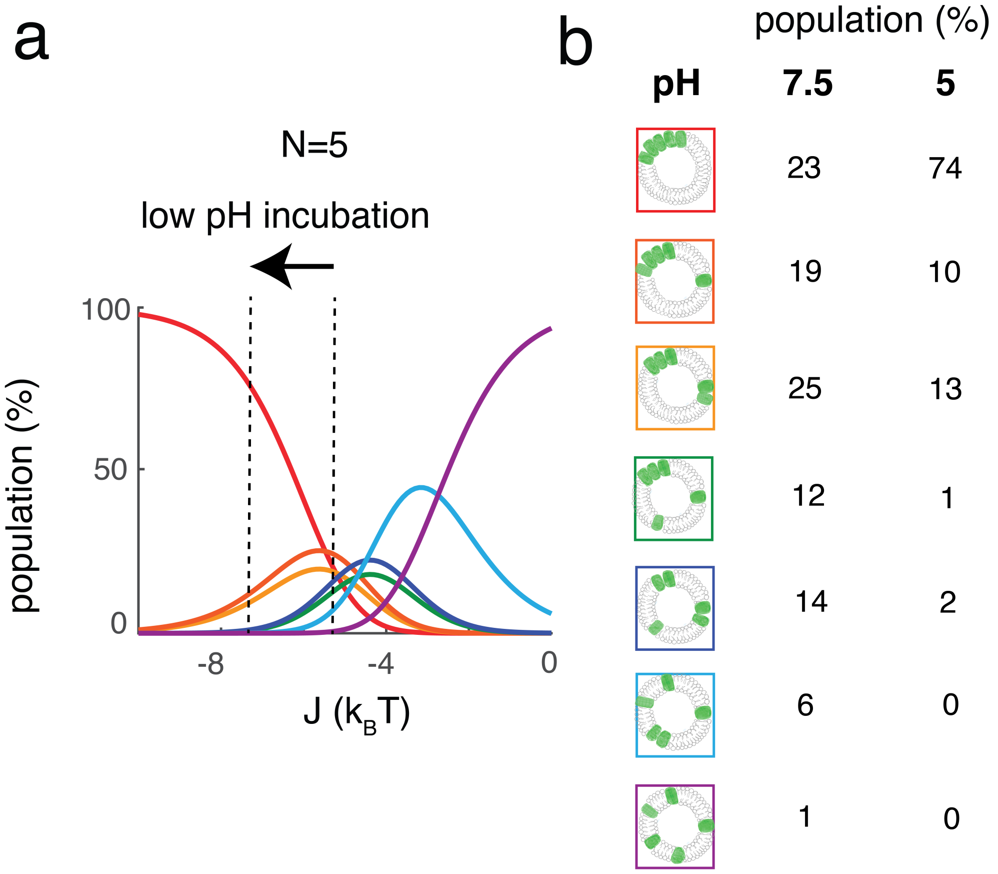

Figure S33: **Shift in population configuration under low pH incubation.** (a) Population of different configurations of the proteoliposome sample with N=5 as a function of LHCII-LHCII interaction energies ($J$). The arrow indicates the shift in $J$ from -5.4 $k_BT$ at pH of 7.5 to -7.2 $k_BT$ at pH 5. (b) Cartoons representing the seven configurations for the N=5 sample and their population percentage under the pH 7.5 and 5. The cartoons of all the configurations are boxed in different colors matched to the population in (a). It is evident from the population percentage that although at neutral pH different configurations with clustered and unclustered proteins coexist, at low pH, the majority of the configurations are heavily clustered. For instance, the population of the configuration where all the proteins are clustered is 23% at pH 7.5. However, the population of this configuration is 74% once incubated at pH 5.

Please note that the first term of the right-hand side of the above equation represents the enthalpy change, and the second term represents the change in entropy for the transition.

Table S9 demonstrates the ΔG, ΔS and ΔS for pH-driven aggregation in samples with <N> of 2,5 and 10. As expected, the aggregation is not entropically favorable. However, the favorable changes in enthalpy lead to the negative ΔGs driving the clustering.

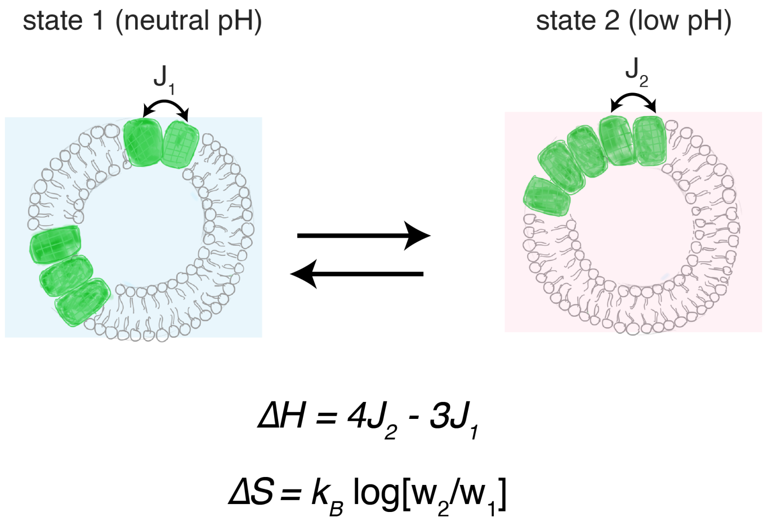

Figure S34: Schematic representation of pH-driven clustering in LHCII-proteoliposome. Based on the relative population of cluster configurations and interaction energies at neutral and low pH, the change in enthalpy (ΔH), entropy (ΔS), and therefore free energy change (ΔG) is calculated.

### 18 Free energy change in light-driven clustering in the chloroplast.

In the previous section, we have discussed various thermodynamic parameters (ΔH, ΔS and ΔG) involving a pH-driven transition from unclustered to a clustered configuration in LHCII proteoliposome. The main difference between a model LHCII proteoliposome system and an in-vivo thylakoid membrane is the protein density. The maximum protein density in the LHCII proteoliposome system studied here is 10 per liposome or 1.3 LHCII per 1000 $nm^2$ of membrane area. On the other hand, the density of LHCII in the thylakoid membrane can be as high as 6.3 per 1000 $nm^2$ of membrane area or ∼5 times higher. This can potentially alter the thermodynamic parameters associated with the quenching in-vivo. To investigate this, we analyzed the freeze-fracture electron micrographs of dark-adapted vio and light-treated zea samples obtained from intact spinach chloroplast [5]. Here, we assume that the vio-dark and zea-light samples represent the systems where LHCII remains in unquenched and quenched configurations, respectively. This is supported by their NPQ values (Table 1 in [5]).

Figure S35a shows the freeze-fracture electron micrographs of vio-dark and zea-light samples. The images are cropped versions of the figure reported in [5]. We used ImagePro software (Media Cybernetics) to analyze these images. First, we identified a few LHCII particles and backgrounds manually from the images. Then these were fed into the 'smart segmentation' feature of the software which thresholds each pixel based on the given intensity values of the particles and the backgrounds. This enables us to segment the image as shown in Figure S35b. The identified areas of the images covered by LHCII particles (green areas in Figure S35) are collected and compared between vio-dark and zea-light samples. As shown in Figure S35c, the median LHCII-covered area of zea-light sample was ∼25% higher compared to vio-dark sample indicating a more clustered configuration of LHCII

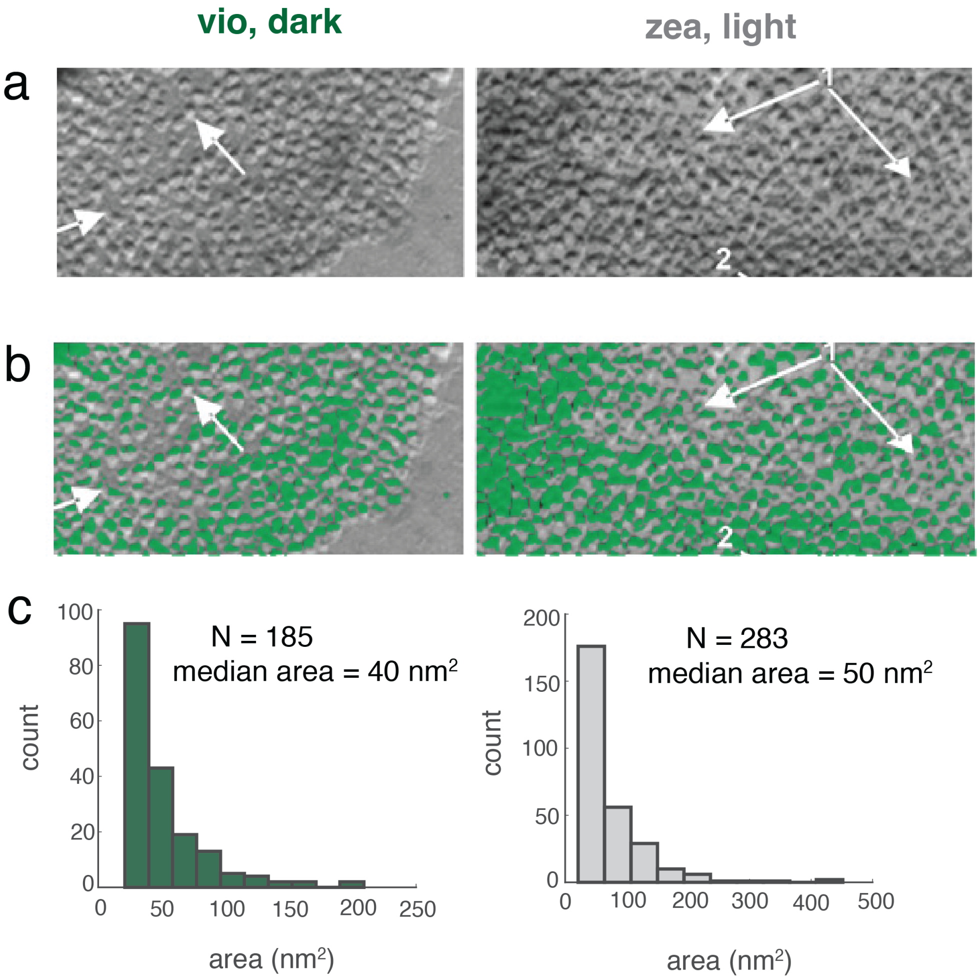

Figure S35: **Image analysis of the freeze-fracture electron micrographs from the spinach chloroplast.**(a) Freeze-fracture electron micrographs of the LHCII periplasmic fracture faces showing dark-adapted vio (left) and light-treated zea (right) from intact spinach chloroplast. This image is obtained from Johnson et al. [5] (Figure 5A, B). The arrows in the original image show the region of enhanced or reduced clustering of LHCII. (b) The original electron micrograph from [5] is cropped and the resulting image is analyzed using Image-Pro software (Media Cybernetics). A few regions of background and particles are identified and fed into the 'smart segmentation' feature of the software to produce a mask that returns the location and size of the LHCII clusters. The masks shown in green are overlayed with the electron micrographs. (c) The distribution of the LHCII cluster area is obtained from the segmented images of dark-adapted vio (left) and light-treated zea (right) samples. The median area of the light-treated zea sample is 25% (40 $nm^2$ vs. 50 $nm^2$) higher compared to the dark-adapted vio sample indicating an overall larger cluster size in the former sample. N = number of clusters in the segmented images.

in the former.

In the next step, we used the segmented images to create particle transforms as shown in Figure S36 a-b. For this, we filled the identified areas with circles that represent LHCII particles. The area of each circle was taken to be 50 $nm^2$ as was observed in [5]. We divided the transforms into regions of $\sim$ 88 $\times$ 88 $nm^2$ square areas which are equivalent to the surface area of proteoliposomes. This enables us to directly compare the ΔH values associated with light-treated clustering of LHCII in chloroplast to pH-induced clustering of LHCII in liposomes. However, we could not extract the ΔS (and therefore ΔG) values from these images using the analytical method described in Sec. 12. The reason for this was the analytical method computes the probability of different configurations only up to $N$ = 20 but the average number of LHCII per 88 nm $\times$ 88 nm square regions in the chloroplast were $\sim$50. Anyway, the change in enthalpy (ΔH) can be measured reliably as this does not involve the probabilities of different configurations involved for a particular $N$. We used a similar methodology to calculate the average enthalpy ($H$) of vio-dark and zea-light samples as was used in the liposome. To give an example, the region highlighted in yellow in Figure S36a contains 45 LHCII proteins. These 45 proteins are in a configuration where they form 19 clusters of 1 (1 $\times$ 19), 6 clusters of two (2 $\times$ 6), two clusters of three (3$\times$ 2), and one cluster of eight (8 $\times$ 1). Therefore, the enthalpy in that region is calculated as:

$$H(J) = (1-1)19J + (2-1)6J + (3-1)2J + (8-1)J = 17J \tag{S43}$$

Similarly, we computed the ΔH for all other square regions. Finally, these mean ΔH values are used to calculate light-induced enthalpy change in the chloroplast in the following way:

$$\Delta H = < H >_{zea} - < H >_{vio} \tag{S44}$$

where $<H>_{vio}$ is the enthalpy averaged over all the 88 nm $\times$ 88 nm regions in vio-dark sample and $<H>_{zea}$ are the enthalpy averaged over all the 88 nm $\times$ 88 nm regions in zea-light sample. Table S9 shows the ΔH for a transition of vio-dark to zea-light configuration of the chloroplast.

Table S9: Thermodynamic parameters for pH-driven aggregation in LHCII-proteoliposomes and chloroplast.

| | <N> | $\Delta H(k_BT)$ | $T\Delta S(k_BT)$ | $\Delta G(k_BT)$ |
|---|---|---|---|---|
| liposome | 2 | -3.6 | -1.6 | -2.0 |
| | 5 | -12.7 | -4.5 | -8.2 |
| | 6 | -15.5 | -5.13 | -10.4 |
| | 10 | -26.2 | -7.2 | -19.0 |
| chloroplast | 50 | -115 | | |

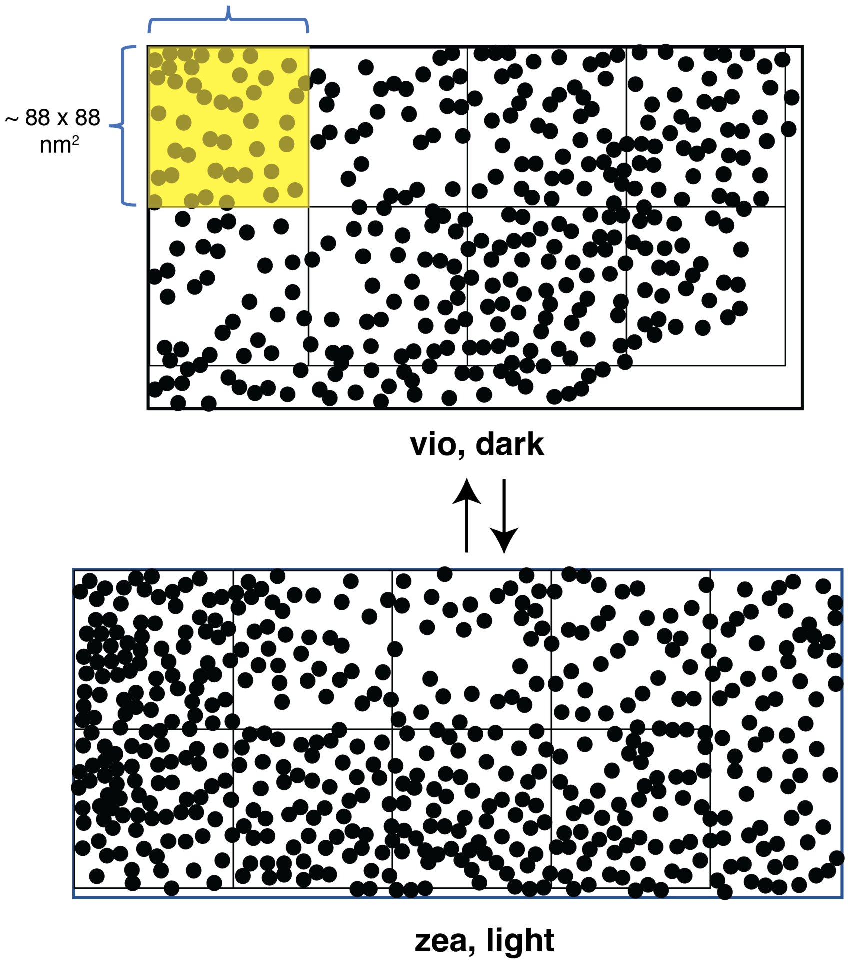

Figure S36: **Free energy in light-driven clustering of LHCII in the chloroplast.** Particle transforms from the segmented images of (a) vio-dark and (b) zea-light samples. The black-filled circles with an area of 50 $nm^2$ represent the LHCII trimers. The images are divided into square regions of $\sim$ 88 nm $\times$ 88 nm to compute relevant thermodynamic parameters, *i.e.*, the change in enthalpy, entropy and free energy. The computations from one such region (highlighted in yellow) are discussed in the text.

### References

[1] Parveen Akhtar et al. "Dependence of chlorophyll fluorescence quenching on the lipid-to-protein ratio in reconstituted light-harvesting complex II membranes containing lipid labels". In: *Chemical Physics* 522 (2019), pp. 242–248.

[2] Emanuela Crisafi and Anjali Pandit. "Disentangling protein and lipid interactions that control a molecular switch in photosynthetic light harvesting". In: *Biochimica et Biophysica Acta (BBA)-Biomembranes* 1859.1 (2017), pp. 40–47.

[3] Vangelis Daskalakis, Sotiris Papadatos, and Ulrich Kleinekathoefer. "Fine tuning of the photosystem II major antenna mobility within the thylakoid membrane of higher plants". In: *Biochimica et Biophysica Acta (BBA)-Biomembranes* 1861.12 (2019), p. 183059.

[4] J Michael Gruber et al. "Singlet–triplet annihilation in single LHCII complexes". In: *Physical Chemistry Chemical Physics* 17.30 (2015), pp. 19844–19853.

[5] Matthew P Johnson et al. "Photoprotective energy dissipation involves the reorganization of photosystem II light-harvesting complexes in the grana membranes of spinach chloroplasts". In: *The Plant Cell* 23.4 (2011), pp. 1468–1479.

[6] Elizabeth Jurrus et al. "Improvements to the APBS biomolecular solvation software suite". In: *Protein Science* 27.1 (2018), pp. 112–128.

[7] Helmut Kirchhoff. "Diffusion of molecules and macromolecules in thylakoid membranes". In: *Biochimica et Biophysica Acta (BBA)-Bioenergetics* 1837.4 (2014), pp. 495–502.

[8] Toru Kondo et al. "Single-molecule spectroscopy of LHCSR1 protein dynamics identifies two distinct states responsible for multi-timescale photosynthetic photoprotection". In: *Nature chemistry* 9.8 (2017), p. 772.

[9] Premashis Manna et al. "Membrane-dependent heterogeneity of LHCII characterized using single-molecule spectroscopy". In: *Biophysical Journal* 120.15 (2021), pp. 3091–3102.

[10] Michael Maus et al. "An experimental comparison of the maximum likelihood estimation and nonlinear least-squares fluorescence lifetime analysis of single molecules". In: *Analytical chemistry* 73.9 (2001), pp. 2078–2086.

[11] Alberto Natali et al. "Light-harvesting complexes (LHCs) cluster spontaneously in membrane environment leading to shortening of their excited state lifetimes". In: *Journal of Biological Chemistry* 291.32 (2016), pp. 16730–16739.

[12] Lauren Nicol and Roberta Croce. "The PsbS protein and low pH are necessary and sufficient to induce quenching in the light-harvesting complex of plants LHCII". In: *Scientific reports* 11.1 (2021), pp. 1–8.

[13] Lauren Nicol and Roberta Croce. "The PsbS protein and low pH are necessary and sufficient to induce quenching in the light-harvesting complex of plants LHCII". In: *Scientific Reports* 11 (1 2021), p. 7415. ISSN: 2045-2322. DOI: 10.1038/s41598-021-86975-9. URL: https://doi.org/10.1038/s41598-021-86975-9.

[14] Mats H M Olsson et al. "PROPKA3: Consistent Treatment of Internal and Surface Residues in Empirical pKa Predictions". In: *Journal of Chemical Theory and Computation* 7 (2 Feb. 2011). doi: 10.1021/ct100578z, pp. 525–537. ISSN: 1549-9618. DOI: 10.1021/ct100578z.

[15] F. Pedregosa et al. "Scikit-learn: Machine Learning in Python". In: *Journal of Machine Learning Research* 12 (2011), pp. 2825–2830.

[16] PG Saffman and M Delbrück. "Brownian motion in biological membranes." In: *Proceedings of the National Academy of Sciences* 72.8 (1975), pp. 3111–3113.

[17] Shreenivas K Sinha. "Classical statistical mechanics of interacting system". In: *Introduction to statistical mechanics*. Alpha Science Int'l Ltd., 2005, pp. 204–239.

[18] Minjung Son et al. "Protein–Protein Interactions Induce pH-Dependent and Zeaxanthin-Independent Photoprotection in the Plant Light-Harvesting Complex, LHCII". In: *Journal of the American Chemical Society* 143.42 (2021), pp. 17577–17586.

[19] Marijonas Tutkus et al. "Aggregation-related quenching of LHCII fluorescence in liposomes revealed by single-molecule spectroscopy". In: *Journal of Photochemistry and Photobiology B: Biology* 218 (2021), p. 112174.

[20] Marijonas Tutkus et al. "Fluorescence Microscopy of Single Liposomes with Incorporated Pigment–Proteins". In: *Langmuir* 34.47 (Nov. 2018), pp. 14410–14418. DOI: 10.1021/acs.langmuir.8b02307. URL: https://doi.org/10.1021/acs.langmuir.8b02307.

[21] Marijonas Tutkus et al. "Fluorescence microscopy of single liposomes with incorporated pigment–proteins". In: *Langmuir* 34.47 (2018), pp. 14410–14418.

[22] Lucas P Watkins and Haw Yang. "Detection of intensity change points in time-resolved single-molecule measurements". In: *The Journal of Physical Chemistry B* 109.1 (2005), pp. 617–628.

[23] Yuri Zaushitsyn et al. "Ultrafast dynamics of singlet-singlet and singlet-triplet exciton annihilation in poly (3-2'-methoxy-5' octylphenyl) thiophene films". In: *Physical Review B* 75.19 (2007), p. 195201.
